# After wounding, a G-protein coupled receptor promotes the restoration of tension in epithelial cells

**DOI:** 10.1101/2023.05.31.543122

**Authors:** Ivy Han, Junmin Hua, James S. White, James T. O’Connor, Lila S. Nassar, Kaden J. Tro, Andrea Page-McCaw, M. Shane Hutson

## Abstract

The maintenance of epithelial barrier function involves cellular tension, with cells pulling on their neighbors to maintain epithelial integrity. Wounding interrupts cellular tension, which may serve as an early signal to initiate epithelial repair. To characterize how wounds alter cellular tension, we used a laser-recoil assay to map cortical tension around wounds in the epithelial monolayer of the *Drosophila* pupal notum. Within a minute of wounding, there was widespread loss of cortical tension along both radial and tangential directions. This tension loss was similar to levels observed with Rok inactivation. Tension was subsequently restored around the wound, first in distal cells and then in proximal cells, reaching the wound margin about 10 minutes after wounding. Restoring tension required the GPCR Mthl10 and the IP3 receptor, indicating the importance of this calcium signaling pathway known to be activated by cellular damage. Tension restoration correlated with an inward-moving contractile wave that has been previously reported; however, the contractile wave itself was not affected by Mthl10 knockdown. These results indicate that cells may transiently increase tension and contract in the absence of Mthl10 signaling, but that pathway is critical for fully resetting baseline epithelial tension after it is disrupted by wounding.

## INTRODUCTION

In response to epithelial damage, an organism must have robust mechanisms for wound repair or else risk exposure to dangers such as infection. Many aspects of the epithelium are damaged by wounding: some cells are lost or die, other cells are damaged but recover, and yet other cells lose cortical tension (O’Connor *et al*., 2021a). In healthy epithelia, tension is maintained by cells pulling on each other through cellular junctions tethered to the actin cytoskeleton. Tension forces are usually balanced, so cells do not move, but temporary adjustments in tension allow epithelial cells to alter their positions in response to the internal or external environment while still maintaining their sheet-like organization (Fernandez-Gonzalez *et al*., 2009; Zulueta-Coarasa and Fernandez-Gonzalez, 2018). By removing cells, wounds impair the ability of pairs of cells to maintain balanced tension. Thus, one aspect of epithelial wounding is a loss of cellular tension, and one aspect of repair is restoring that tension.

A wound-induced change in tension has been considered as a possible early wound signal, sensed by surrounding epithelial cells, triggering wound-repair behaviors (Xu and Chisholm, 2011; Antunes *et al*., 2013; Enyedi and Niethammer, 2015; Cao *et al*., 2017; Franco *et al*., 2019); however, the extent of this potential signal around wounds has not yet been explored. For example, how far from the wound do changes in tension extend? Do tension levels change in a gradient from the wound? Further, specific mechanisms to restore cellular tension after wounding have not been identified. Here we answer these questions in the particular case of the *Drosophila* pupal notum.

The pupal notum epidermis is an epithelial monolayer that offers an excellent system to probe the dynamics of wound repair. This tissue can be imaged *in vivo* continuously – before, during, and after laser ablation, throughout the entire process of wound repair – and pupae can survive wounding and imaging to eclose and walk away (Shannon *et al*., 2017). Using this system and harnessing the power of fly genetics, our laboratories previously identified a wound-induced signaling pathway that acts through the cell surface receptor Mthl10, a G-protein coupled receptor (GPCR) coupled to G_αq_. The wound-induced activation of Mthl10 is transduced downstream by G_αq_ and PLCβ, generating IP_3_ to release Ca^2+^ from internal stores through the IP_3_ receptor (IP3R; O’Connor *et al*., 2021b). Although Mthl10 improves viability after wounding, how its activation affects cells and promotes wound repair is not known. Two potential clues are that the second messenger downstream of Mthl10, i.e., Ca^2+^, has a profound impact on the actin cytoskeleton (Hunter *et al*., 2015; Wales *et al*., 2016; Ready and Chang, 2021), and it has been shown that increasing Ca^2+^ alone is sufficient to induce cellular contraction (Kong *et al*., 2019).

In this study, we investigated wound-induced changes in cellular tension. Using laser-induced recoil to infer cellular tensions quantitatively (Ma *et al*., 2009), we mapped changes in tension at different locations and time points after wounding. We found that that within one minute, tension was lost around the wound in a widespread and uniform manner, in both the radial and tangential directions. Tension was restored over the next ∼10 minutes in a spatiotemporal pattern, from outward in, as a wound-induced contractile wave advanced toward the wound. Restoring tension required signaling by Mthl10 and its downstream release of calcium via the IP3R. Even when wounds were administered using a method that did not generate an evident contractile wave, *mthl10* was still required for the restoration of tension, suggesting that Mthl10 may restore tension around many types of epithelial wounds.

## RESULTS

### Laser Recoil Measures Changes in Tension across the *Drosophila* Notum

An epithelial sheet consists of cells that are connected with each other under tension. Breaking those connections by creating a wound in the tissue would be expected to lead to changes in tension. To probe cortical tension around a wound, we used a laser-induced recoil assay. This assay is performed by severing an apical epithelial cell border with laser micro-ablation and measuring the retraction velocity of the corresponding tricellular junctions at the ends of the severed border; higher recoil velocities correspond to higher cortical tension, while lower recoil velocities correspond to lower cortical tension (Fig. 1A-D, Movie 1). We measured tension at specific sites, using the border of the *pnr* domain as a landmark (Fig. 1A,B). In control samples, the tissues on each side of the notum’s *pnr*-domain boundary have no discernable difference in cell morphology, Ecad-GFP or MyoII-GFP fluorescence (Fig. S1A-B). At each site on both sides of the boundary, we measured tension in two axes, severing cell borders that were aligned with the anterior-posterior (AP) and the mediolateral (ML) axes of the tissue (Fig. 1C). To establish the assay and set a baseline, we measured tension in unwounded control samples to map epithelial tension across the notum. We found that tension was fairly constant at all locations tested for both ML and AP epithelial cell borders, with no evidence of location-specific or *pnr*-specific effects (Fig. 1E). This finding indicates that epithelial tension in the unwounded notum is reasonably constant.

**Figure 1.**
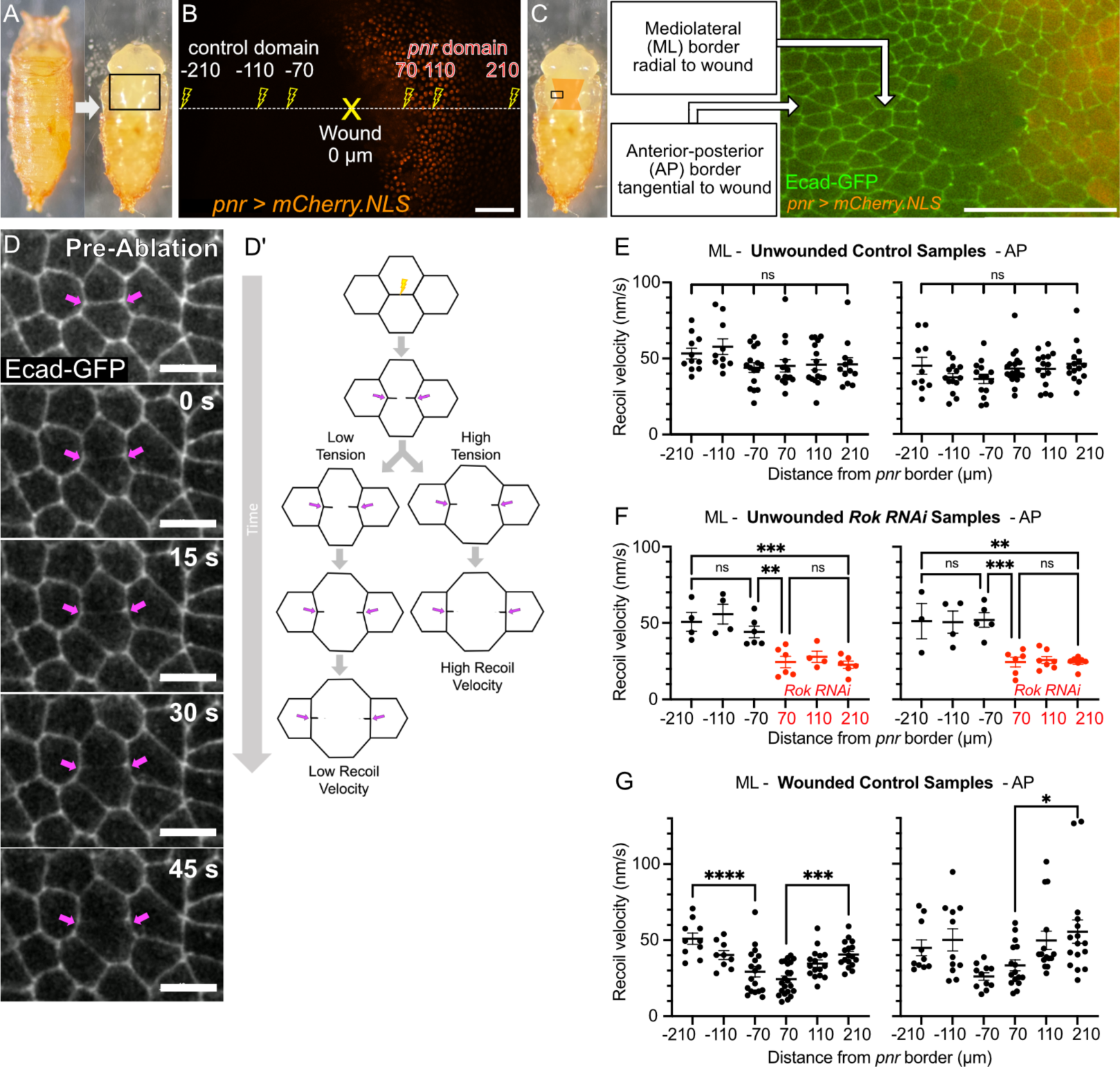
Tension is lost around wounds. **(A)** *Drosophila melanogaster* pupa (left) with pupal case removed (right) to expose notum (outlined by black rectangle). **(B)** Map of distances where tension was measured. Wound is designated as 0 µm (yellow X). Tension was measured at ∼70, 110, and 210 µm on either side of the wound (yellow lightning bolts). Map is overlaid on an image of the notum with *pnr* domain labeled by red nuclei. Closest distance (∼70 µm) corresponds to the edge of nuclear membrane damage visible in wounded samples. **(C)** Two types of borders measured: mediolateral (ML) borders, which are radial to the wound at this location, and anterior-posterior (AP) borders, which are tangential to the wound at this location. **(D)** Example of laser-induced recoil for a mediolateral cell border. Tricellular junctions of ablated cell border indicated by magenta arrows. See also: Movie 1. **(D′)** Diagram illustrating how laser-induced recoil reports cortical tension: low tension corresponds to low recoil velocity; high tension corresponds to high recoil velocity. **(E)** In unwounded control samples, tension was fairly constant at all locations tested for both ML and AP borders. *n* = 10 to 16 for each bin. **(F)** In unwounded *pnr > Rok RNAi* samples, significantly lower tension was detected in both ML and AP borders. Reduced tension was confined to the domain expressing *Rok RNAi*. *n* = 3 to 7 for each bin. **(G)** In wounded control samples, tension was reduced in a gradient for both ML and AP borders, with lower tension closer to the wound. Measurements taken 1-10 minutes post-wound. *n* = 9 to 22 for each bin. Graph bars represent mean ± SEM. *p<0.05, **p<0.01, ***p<0.001, ****p<0.0001 by one-way ANOVA with multiple comparisons. Scale bars = 50 µm (B and C), or 10 µm (D).

Because there are no *pnr*-domain-specific effects in control pupae, we can drive genetic manipulations within the *pnr* domain using *pnr-Gal4*, with the neighboring domain acting as an internal control. It was unclear, however, whether a loss of tension generated in the *pnr* domain would equilibrate across the notum to the control domain or whether it would remain in the *pnr* domain. To determine if we could detect different levels of tension across the notum, we mapped tension in unwounded samples with *Rok RNAi* expressed in just the *pnr* domain. Rok, or Rho kinase, modulates stress and contractility by phosphorylating cytoskeletal proteins including myosin II, and inhibiting or knocking down *Rok* reduces cortical tension (Mizuno *et al*., 1999). Even with this domain-specific knock down of *Rok*, there was no discernable difference in cell morphology or Ecad-GFP fluorescence between the two domains (Fig. S1C). Interestingly, there was a decrease in cortical tension at locations in the *pnr* domain with *Rok RNAi* compared to the control domain (Fig. 1F). This trend was clear for both ML and AP borders. Tension in the control domain was relatively constant and similar to previous levels; tension in the *pnr>Rok RNAi* domain was also relatively constant, but at a decreased value.

These results demonstrate that although the notum is a continuous epithelium, domain-specific differences in cortical tension can exist and can be detected. To better understand how a continuous tissue could maintain regions with different cortical tension, we ran vertex-model simulations (Fletcher *et al*., 2014) of a tissue in which one side of the tissue patch had cells with reduced contractility. We chose model parameters similar to those from Tetley *et al*. (2019), but with contractility reduced by 63% on one side. These simulated tissues reach mechanical equilibrium (Fig. S1D) and yield simulated recoil velocity patterns that match experiments: recoils are approximately half as fast (59%) in the domain with reduced contractility. A close look at the modeled forces shows that equilibrium is attained when the mesoscopic tensile stress, i.e., that averaged over several cells, is the same on both sides. Nonetheless, each domain generates that tensile stress differently. In the domain with reduced contractility, the vertex model’s perimeter terms contribute less, and its cells stretch a little until the area-based tensions increase to compensate. In the idealization of a vertex model, this increase in cell area is measurable and significant, but it is small (3%; inset to Fig. S1D) and would be dwarfed by the natural cell-to-cell variation in a living epithelium. Cells within the notum may use similar or even additional compensation, e.g., variation in the strength of their attachments to the cuticle and/or basement membrane. Even when cortical tension is reduced in some cells, an epithelium has several parallel means by which it can reach a given mesoscopic tension.

### Wounding Reduces Cortical Tension Locally

Next, we measured cortical tension around wounds. Wounds were generated using the same laser used for measuring recoil velocity but at 4-fold higher power, and they were targeted at the boundary of the mCherry.NLS-labeled *pnr* domain. Previous work by our group found that this type of laser ablation generates specific and reproducible types of cellular damage at different distances from the center of the wound (O’Connor *et al*., 2021a). One of the most far-reaching types of damage is the loss of nuclear membrane integrity, visible in the *pnr* domain by the loss of mCherry.NLS from the nuclei, extending out ∼70 µm from the center of the wound. We used this nuclear membrane damage as the landmark for the most proximal testing location, measuring tension at the edge of this visible region of damage, and at two more fixed distances distal to this location on the *pnr* side. We mirrored these three locations on the other side of the *pnr* border, testing tension at the same three radii in the control domain. For these locations, ML borders were radial to the wound and AP borders were tangential to the wound. A few minutes after wounding control samples, tension was reduced in a gradient on both sides of the *pnr* domain boundary, with the lowest tension found closest to the wound (Fig. 1G). This lowest tension close to the wound was similar to the level observed with *Rok RNAi*, and the highest tension at the furthest locations, ∼210 μm from the wound, was similar to that of unwounded samples. The gradient was most clearly significant for ML cell borders. There was a similar trend for AP borders, but those recoil measurements were more variable and on the edge of significance. What we did not observe was the transfer of tension from radial borders (ML) to tangential ones (AP) that one would expect around a hole in an elastic sheet (Timoshenko and Goodier, 1951; Vishay Micro-Measurements, 2007). Instead, within a few minutes after wounding, tension in apical epithelial cell borders was reduced at points closer to the wound.

### Tension is Restored over Time

After generating a spatial map of tension around wounds, we generated a temporal map, investigating if post-wounding tension at a certain location changed over time. Our previous measurements were taken between 1 to 10 minutes after wounding. To create our temporal map, we measured tension at ∼70 µm at 1-2 minutes and 15-18 minutes after wounding using mediolateral borders. We observed a dramatic restoration of tension to pre-wound levels at the later times, indicating that tension is dynamically regulated around wounds (Fig. 2A,B).

**Figure 2.**
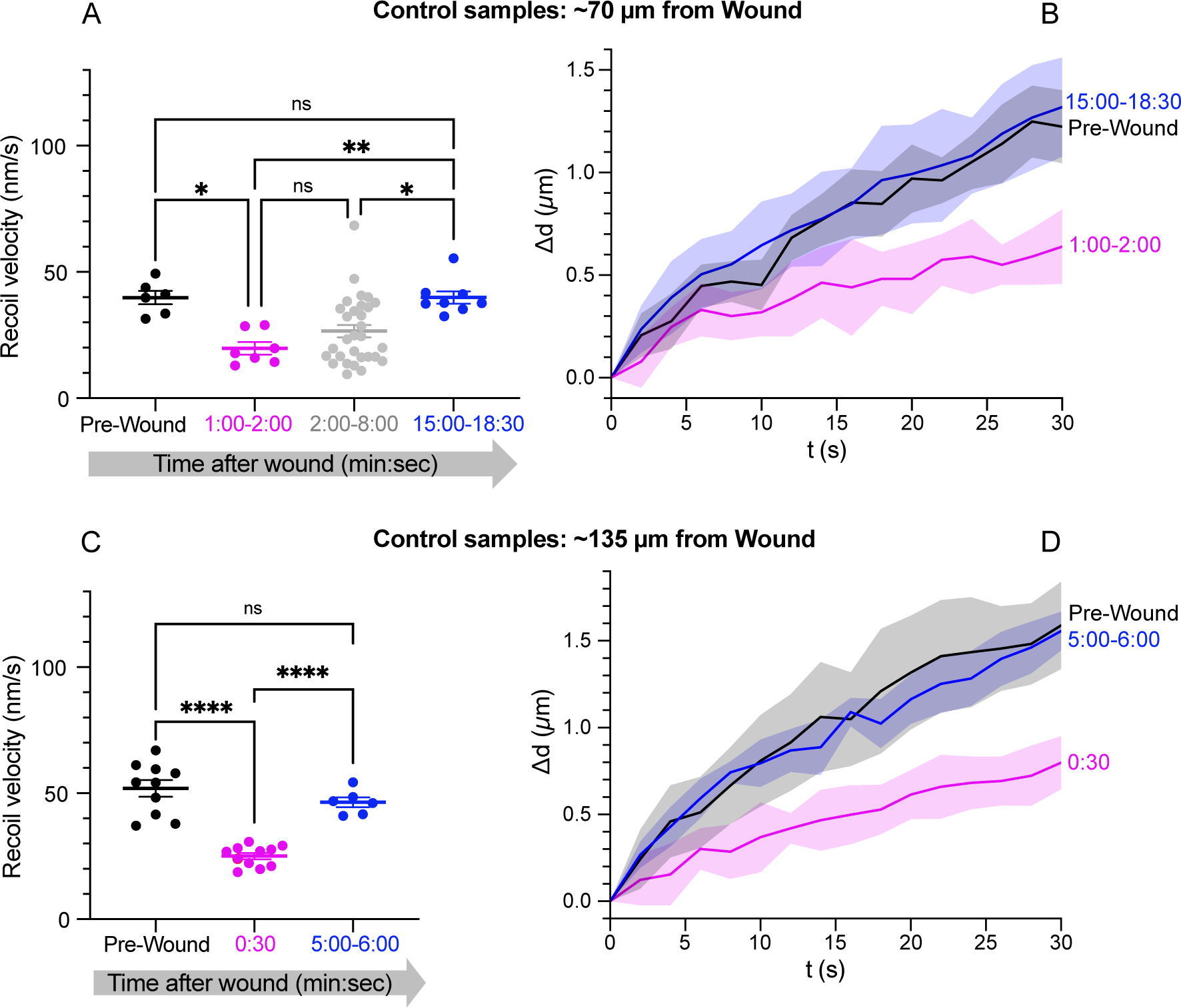
Tension is restored over time. **(A)** At ∼70 µm from the wound, cortical tension is reduced by half by 1-2 min after wounding and restored to pre-wound levels within 15-18 min. Data collected 2-8 min after wounding shows a mix of tension levels (light gray, from Fig. 1G); *n* = 6, 7, 30 and 9. **(B)** Data from panel A showing average recoil displacements versus time. **(C)** At ∼135 µm from the wound, cortical tension is reduced by half by 30 sec after wounding and restored to pre-wound levels within 5-6 min; *n* = 10, 11 and 6. **(D)** Data from panel C showing average recoil displacements versus time. Graph bars represent mean ± SEM. *p<0.05, **p<0.01, ****p<0.0001 by one-way ANOVA with multiple comparisons. Shaded regions in B and D represent ± one standard deviation. All measurements conducted on ML borders.

Given these results, we realized the observed tension gradient around wounds had a temporal component, so we investigated if tension at distal locations would be reduced immediately (< 1 minute) after wounding. We measured tension in ML borders at ∼135 µm from the wound about 30 seconds after wounding and compared it to tension at the same distance and orientation 5-6 minutes after wounding. Excitingly, at the 30 second time point, we observed an initial loss of tension at ∼135 µm to levels similar to *Rok RNAi*, and tension was restored to pre-wound levels at this location by 5-6 minutes (Fig. 2C,D). Our previous measurements over a wider and slower time range (Fig. 1G) had missed this initial decrease in tension farther from wounds because we measured it after it was restored. Thus, cortical tension was restored in a temporal and spatial gradient – first distally, then proximally.

### A Contractile Wave Moves toward the Wound

A previous study reported another distal-to-proximal phenomenon after laser wounding: a symmetrical wave of apical contractions that starts as a distal ring within ∼2 minutes after wounding and then travels inward to the wound margin (Antunes *et al*., 2013). The apical surfaces of the outer ring of cells contract and expand synchronously over ∼5 minutes; as the cells of that ring expand, their proximal neighbors begin contracting. This pattern propagates inward across 4-5 rows of cells until the contracting ring reaches the cells at the leading edge of the wound ∼12-14 minutes after wounding (Fig. 3A, Movie 2). Each cellular contraction is preceded by a temporary apical accumulation of actin and myosin (Antunes *et al*., 2013; Fig. 3A-C, Movie 3). Given the involvement of actin and myosin, we wondered if the contractile wave might be related to tension restoration around the wound.

**Figure 3.**
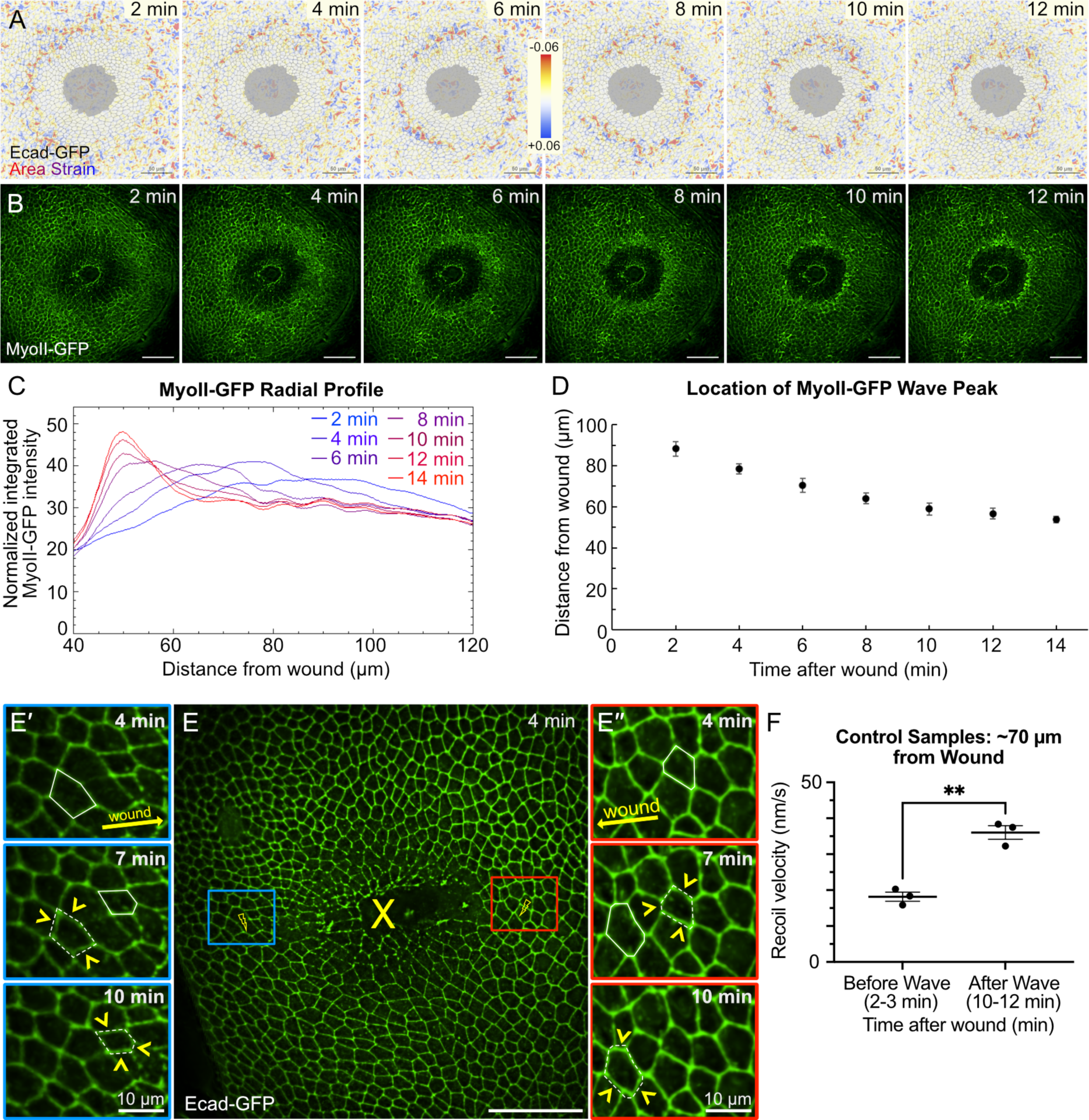
Tension is restored as the contractile wave moves toward the wound. **(A)** A wave of cell contractions travels inward over 12 minutes after wounding and can be visualized in Ecad-GFP-labeled pupae by measuring displacement fields between successive frames and calculating the local area strain (one of *n* = 5 shown). Negative values correspond to contractions. See also Movie 2. **(B)** The contractile wave can also be visualized as a wave of apical MyoII-GFP accumulation traveling inward after wounding (one of *n* = 7 shown). See also Movie 3. Note that A,B show different wounds. **(C)** Peaks of radial MyoII-GFP intensity from B move closer to the wound over time, tracking progression of the contractile wave. **(D)** The average location of the wave over time, as determined from data similar to panel C averaged over *n* = 7 wounds. **(E-E′′)** Using Ecad-GFP to monitor the contractile wave, tension was measured before (blue) and after (red) the contractile wave passed through a cell ∼70 µm from the wound. Yellow arrowheads in E′ and E′′ show contracting cells. See also Movie 4. **(F)** Tension at ∼70 µm from the wound is restored as the wave passes through. Graph bars represent mean ± SEM. *n* = 3 and 3; **p<0.01 by paired sample t-test. Scale bars = 50 µm (A, B, and E), or 10 µm (E′ and E′′).

The tension measurements above indicate that cells at ∼70 µm from the wound have reduced tension at 2 minutes after wounding and restored tension after 15 minutes. These two times bracket the expected passage of the propagating contractile wave (Fig. 3D). To more closely assess the correlation, we imaged the wave by tracking cell contraction and measured tension at ∼70 µm from the wound both before and after directly observing the contractile wave passing through that location (Fig. 3E-E′′, Movie 4). To prevent the first measurement from interfering with the second, the paired measurements were acquired on different sides of the wound. We found that tension at this location was low before the wave passed through and was then restored to near pre-wound levels shortly after the wave passed through (Fig. 3F). Thus, we conclude that the restoration of tension is temporally correlated with the contractile wave; however, as shown below, physical and genetic manipulations can separate the two phenomena.

### Mthl10 is Required to Restore Tension

Our labs previously discovered that the GPCR Methuselah-like 10 (Mthl10) is activated in epithelial cells surrounding a wound between 45-75 seconds after wounding (Shannon *et al*., 2017; O’Connor *et al*., 2021b). The GPCR triggers the G_αq_-signaling pathway, resulting in the release of calcium via the IP_3_ Receptor (IP3R) in the endoplasmic reticulum.

To investigate whether Mthl10 signaling modulates tension, we measured tension in both unwounded and wounded samples with *mthl10 RNAi* expressed in the *pnr* domain to knock down Mthl10. Similar to previous unwounded control samples, unwounded tension was relatively homogeneous across both control and knockdown regions for both ML and AP cell borders (Fig. 4A). In the wounded samples, tension on the control side followed a gradient like that before, with reduced tension at points closer to the wound – an effect that the experiments in Figs. 2 and 3 showed was due to a time- and distance-dependent restoration of tension after wounding. On the knockdown side, post-wound tension fell and remained uniformly low at all distances tested (Fig. 4B), suggesting that Mthl10 is required for restoring tension. These results were confirmed independently with a second *UAS-mthl10 RNAi #2* line (Fig. S2A). To test if Mthl10 signaling restores tension through IP3R, we knocked down *IP3R* and found that it was required to restore tension, indicating the release of calcium from the ER restores tension around wounds (Fig. 4C). We also tested an additional far distal location in the *pnr > IP3R* domain at ∼270 µm from the wound and found unwounded levels of tension, indicating that wounding reduces tension in a limited region around the wound (Fig. S2B).

**Figure 4.**
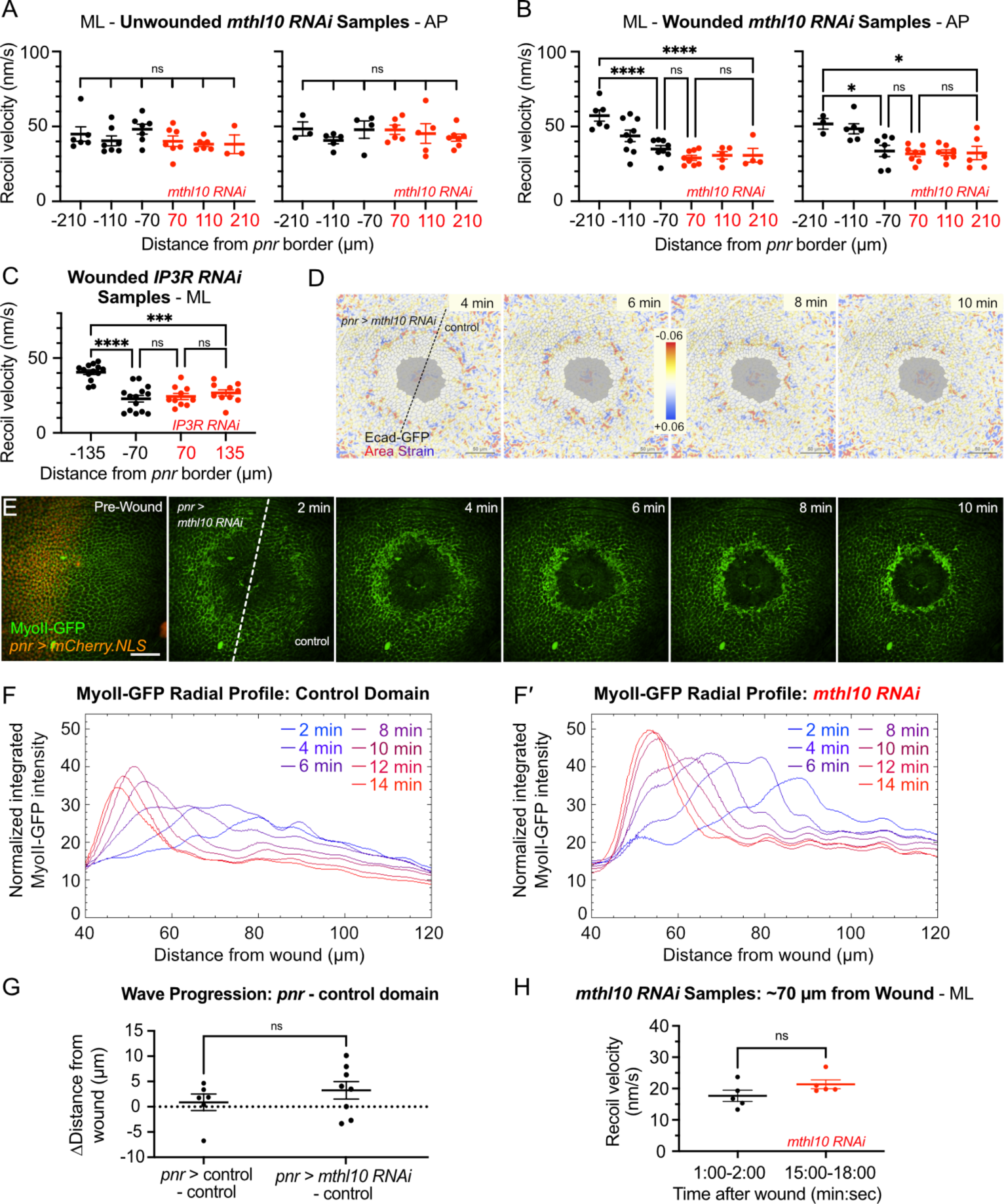
The restoration of tension after wounding requires the GPCR Mthl10. **(A)** In unwounded flies, knockdown of *mthl10* within the *pnr* domain had no effect on cortical tension for both ML and AP borders; *n* = 3 to 7 per bin. **(B)** The wound-induced loss of tension persists after wounding in the *pnr* domain where *mthl10* was knocked down, indicating that Mthl10 is required for the restoration of tension; *n* = 3 to 9 per bin. **(C)** IP3R is downstream of Mthl10 and is also required for the restoration of tension after wounding, indicating that Mthl10 signaling restores tension through IP3R. Measurements in B-C taken 1-10 minutes post-wound; *n* = 10 to 14 per bin. **(D-G)** The contractile wave does not require *mthl10.* Wave symmetry was not disturbed when *mthl10* was knocked down in the *pnr* domain, as visualized with area strain maps calculated for displacement fields between successive Ecad-GFP images (D; *n* = 4; see also Movie 5) or with MyoII-GFP (E; *n* = 7, see also Movie 6). Radial profile analysis (F and F′) showed no significant difference between control and *pnr* domains in *pnr > mthl10 RNAi* samples compared to control samples (G; *n* = 6 and 8). **(H)** The contractile wave exists without *mthl10*, but it requires *mthl10* to restore tension after wounding. Tension is not restored at ∼70 µm even 15 minutes post-wound in *pnr > mthl10RNAi* samples. Graph bars represent mean ± SEM; *n* = 5 and 5; *p<0.05, ***p<0.001, ****p<0.0001 by one-way ANOVA with multiple comparisons (A, B, C), unpaired t-test (G), or paired sample t-test (H). Scale bars = 50 µm.

Surprisingly, knockdown of *mthl10* did not disturb the rate of wound closure (Fig. S3), the formation of syncytia around the wound (Fig. S3; White *et al*., 2023), nor the contractile wave (Fig. 4D-G; Movies 5 and 6). The wave was visible as it traveled across cell borders toward the wound, even in the absence of Mthl10 (Fig. 4D), and the pattern of MyoII-GFP levels moving toward the wound was also similar between control and knockdown regions around the wound (Fig. 4E-F’). To see if the rate of travel was altered by *mthl10* knockdown, we analyzed the location of the wave on the two sides of the wound (control and *pnr*) in both control samples (+ and +) and in samples in which *mthl10* was knocked down in the *pnr* domain (+ and *mthl10 RNAi*). We found that in both cases, the wave was at similar locations on the two sides of the wound, indicating that Mthl10 did not significantly alter the wave’s rate of travel (Fig. 4G). Finally, when we knocked down *mthl10*, tension remained low over the first 15 minutes at ∼70 µm from the wound (Fig. 4H) in contrast with the control data from Figure 2A. Thus, the correlation between the contractile wave and the restoration of tension is broken in the absence of Mthl10.

### Scanned Wounds Also Require Mthl10 to Restore Wound-Induced Tension Loss

The laser wounds we analyzed thus far generate several different types of cellular damage arrayed in a regular pattern around the center of the wound (O’Connor *et al*., 2021a). To ask if these damaged cells contribute to the loss of tension around wounds, we measured tension around a different type of wound, scanning ablation, in which low-energy laser pulses are scanned repeatedly across a circular area to lyse all cells. This process takes about 2 minutes to scan an area matching the area of cell death in the wounds above. To confirm that these repeated low-energy laser pulses were killing cells and not just photobleaching them, we conducted scanning ablation on a notum where cell borders were labeled with fluorescently tagged p120-catenin (p120ctn-RFP) and chromatin was labeled with His2AvGFP. The results of this experiment are shown in Fig. S4A. The scanning ablation process destroys cell borders and nuclei but leaves behind lots of still-fluorescent fragmented chromatin (histones) that is not motile (observed at 0-2h). This debris is eventually cleared from the wounded region (3-5 h), presumably by circulating macrophage-like hemocytes. Fluorescence from neither marker recovers until the scanned-ablation region is repopulated by nuclei moving in from outside the exposed region.

These wounds showed no evidence of a contractile wave (Fig. 5A,B; Fig. S4; Movies 7 and 8). Further, within a few minutes of completing scanning ablation in control samples, measured tension levels across the notum were similar to pre-wound tensions (Fig. 5C), suggesting either that tension was never lost or that it was completely restored before our measurements. To distinguish these possibilities, we generated scanned wounds in pupae with *mthl10* knocked down in the *pnr* domain and found that tension levels in the *pnr* domain were reduced to levels similar to the *Rok* knockdown (Fig. 5D). We conclude that tension was lost temporarily around scanned wounds, but quickly restored by Mthl10. These results suggest that tension loss and restoration is a general phenomenon of epithelial wounds.

**Figure 5.**
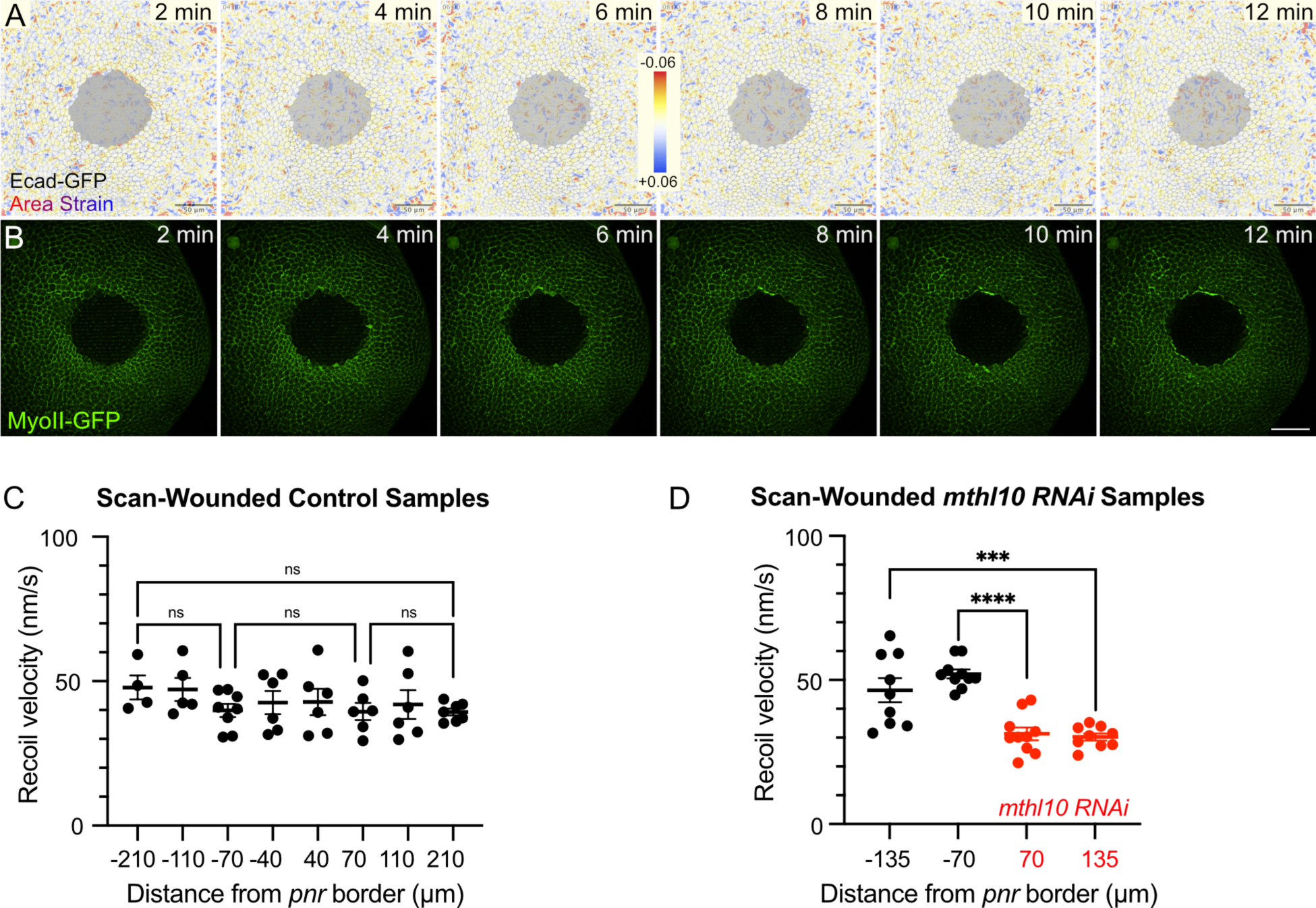
Mthl10 is also required for tension restoration around scanned wounds. **(A-B)** In scan-wounded control samples, there is no evident wave of contraction (*n* = 5; see also Movie 7) or myosin accumulation (*n* = 5; see also Movie 8). Different wounds are shown in A,B. **(C)** Within ∼2-10 minutes of completing a scanned wound, post-wound tension is fairly uniform at all locations tested; *n* = 4 to 8 per bin. **(D)** In contrast, for scan-wounded *pnr > mthl10 RNAi* samples, there is reduced tension in the knockdown domain, indicating that scanned wounds also require Mthl10 for tension restoration; *n* = 9 to 10 per bin. All measurements were taken on ML cell-cell borders. Graph bars represent mean ± SEM. ***p<0.001, ****p<0.0001 by one-way ANOVA with multiple comparisons. Scale bars = 50 µm.

## DISCUSSION

In this study, we used a laser-induced recoil assay to measure changes in cortical tension after epithelial wounding in the *Drosophila* notum. Although before wounding, epithelial cells have relatively constant cortical tension, we find that wounds trigger loss of tension across a discrete area. Within this area, tension is reduced uniformly, to similar levels as when *Rok* is knocked down, and to similar extents along orthogonal axes. Tension is then restored over 2-10 minutes – first at locations distal to the wound and proceeding proximally. This restoration of tension requires calcium signaling through the IP_3_ Receptor downstream of the wound-responsive G-protein-coupled receptor Methuselah-like 10 (Mthl10). Our labs previously found that *mthl10* knockdown throughout the notum decreased pupal survival by ∼30% (O’Connor *et al*., 2021b). Interestingly, this decreased survival does not arise from differences in wound closure rates (Fig. S3). The importance of Mthl10 for wound survivability may instead be due to the newly discovered tension restoration function of Mthl10 signaling.

The term “tension” is used to represent multiple different phenomena in mechanobiology. For the laser-recoil assays used here, the measured tension is a combination of membrane tension, cell-cell adhesive zippering, and actomyosin activity – all together contributing to cortical tension. Variability in the measurements arises from both the stochastic nature of laser ablation and cell-to-cell variability in the epithelium: each measured point averages tension over a local region of four cells (see Fig. 1D′). The local nature of these measurements is illustrated by the experiments with Rok knockdowns. When the knock down is conducted in a limited region (*pnr* domain), measured tension is lower throughout that domain, and yet tension is normal as close as 70 µm (∼7 cell diameters) outside this region. Although mechanical effects can act over long distances in an epithelium (Alisafaei *et al*., 2021), these experiments show that local regions can maintain significantly higher or lower cortical tension. There are multiple compensatory mechanisms by which adjacent domains in an epithelium can have different cortical tension and nonetheless maintain balanced mesoscopic tensions, e.g., through redistribution of the tension from cell-cell interfaces to medial cell regions (Fig. S1D) or attachment to extracellular matrix.

Before wounding, cortical tensions in control epithelia were relatively uniform and isotropic, i.e., having no preferred direction. Our naive expectation was that making a circular wound would yield a pattern of tension changes matching those around a hole in a viscoelastic sheet. This pattern would have two orthogonal gradients: tensions in radial directions would be reduced around the wound and then gradually rise toward the uniform far-field tension; those in tangential directions would be stronger circling the wound and gradually fall toward the far-field limit (Timoshenko and Goodier, 1951; Vishay Micro-Measurements, 2007). Instead, the measurement showed that tension was reduced to a similar extent in both radial and tangential directions. Although our initial experiments appeared to uncover a spatial gradient of tension loss (e.g., Fig. 1G), further investigation revealed that this result was actually a spatiotemporal gradient of tension restoration. When tension levels were analyzed in the *Mthl10* knockdowns, which were unable to restore tension, we found that wounding induced a loss of tension uniformly out to at least 210 µm from the wound center (although not as far as 270 µm; Fig S2B). Measurements in control pupae as fast as 30 s after wounding confirmed this uniform loss out to at least 135 µm, and its distal to proximal restoration over the next ∼10 minutes.

What could explain this widespread, isotropic and uniform loss of tension immediately after wounding? The low-tension region extends considerably farther than the area of plasma membrane damage caused by the laser-induced cavitation bubble (∼ 110 µm; Shannon *et al*., 2017), indicating that tension loss is not an artifact of cavitation; further, scanned wounds with much smaller cavitation bubbles still lose tension, discernible in *Mthl10* knockdowns, indicating that tension loss is independent of cavitation and plasma membrane damage. One potential explanation for the uniform widespread loss of tension is that cellular mechanics are both passively and actively nonlinear. This nonlinearity around wounds may be adaptive, as tension-induced recoil after wounding would increase the size of the wound; reducing tension may thus limit further damage. An analogy for this possibility would be when clothing gets snagged on a thorn – pulling will rip the clothing but relaxing (i.e., reducing tension) prevents ripping.

The widespread and uniform loss of tension around wounds is fast, occurring within 30 sec (the limit of our ability to make a wound and retarget the laser to measure tension). Tension is then restored over the next ∼ 10 min in a distinct spatiotemporal pattern, from outward in. Tension restoration requires the cell surface G-protein coupled receptor Mthl10, which has fast activation kinetics starting ∼45 sec after wounding, and a distinct spatial domain of activation around wounds (O’Connor *et al*., 2021b), both of which are consistent with the spatiotemporal pattern of restoring cellular tension. However, there is one important difference: Mthl10 activation starts near the wound and moves outward, but tension restoration starts at the edge of the low-tension domain and moves inward. Although there are many possible explanations for this outward-in phenomenon, the simplest might be that cells farther from the wound are less damaged than those closer in (O’Connor *et al*., 2021a), and so they are able to restore tension sooner. This attractive hypothesis is at present untestable: we could reduce the damage gradients by creating scanned ablation wounds, but creating these requires ∼ 2 min, preventing us from measuring dynamic changes in tension on few-minute time scales. Another explanation for the outward-in restoration of tension is that cells must pull on fully-tensioned neighbors to restore their own tension, available only to the outer ring of low-tension cells.

Another phenomenon that travels distal to proximal around wounds is a contractile wave, previously reported by Antunes *et al*. (2013). Coordination between the contractile wave and the restoration of tension is summarized in Figure 6. The wave first becomes evident when a ring of cells ∼100 µm from the wound simultaneously contract. As these cells then relax and expand, their wound-proximal neighbors contract, and the process repeats over about 10 minutes until the contractile wave meets the wound margin. The spatiotemporal dynamics of this wave seem to match the Mthl10-mediated restoration of tension, and indeed we found that tension is low before the wave and is restored after it passes through. Interestingly, though, the wave continues unperturbed in *mthl10* knockdown cells. Propagation of the wave requires cells to increase tension as they contract, and Antunes *et al*. (2013) noted that these contracting cells also transiently accumulate more actin and myosin at their apical surfaces, so *mthl10* knockdown cells must be able to temporarily increase cellular tension; however, they are unable to maintain this tension after the wave passes through. We thus propose that Mthl10 regulates a ratchet-like mechanism, maintaining high tension after the contractile wave temporarily restores it through cellular contraction.

**Figure 6.**
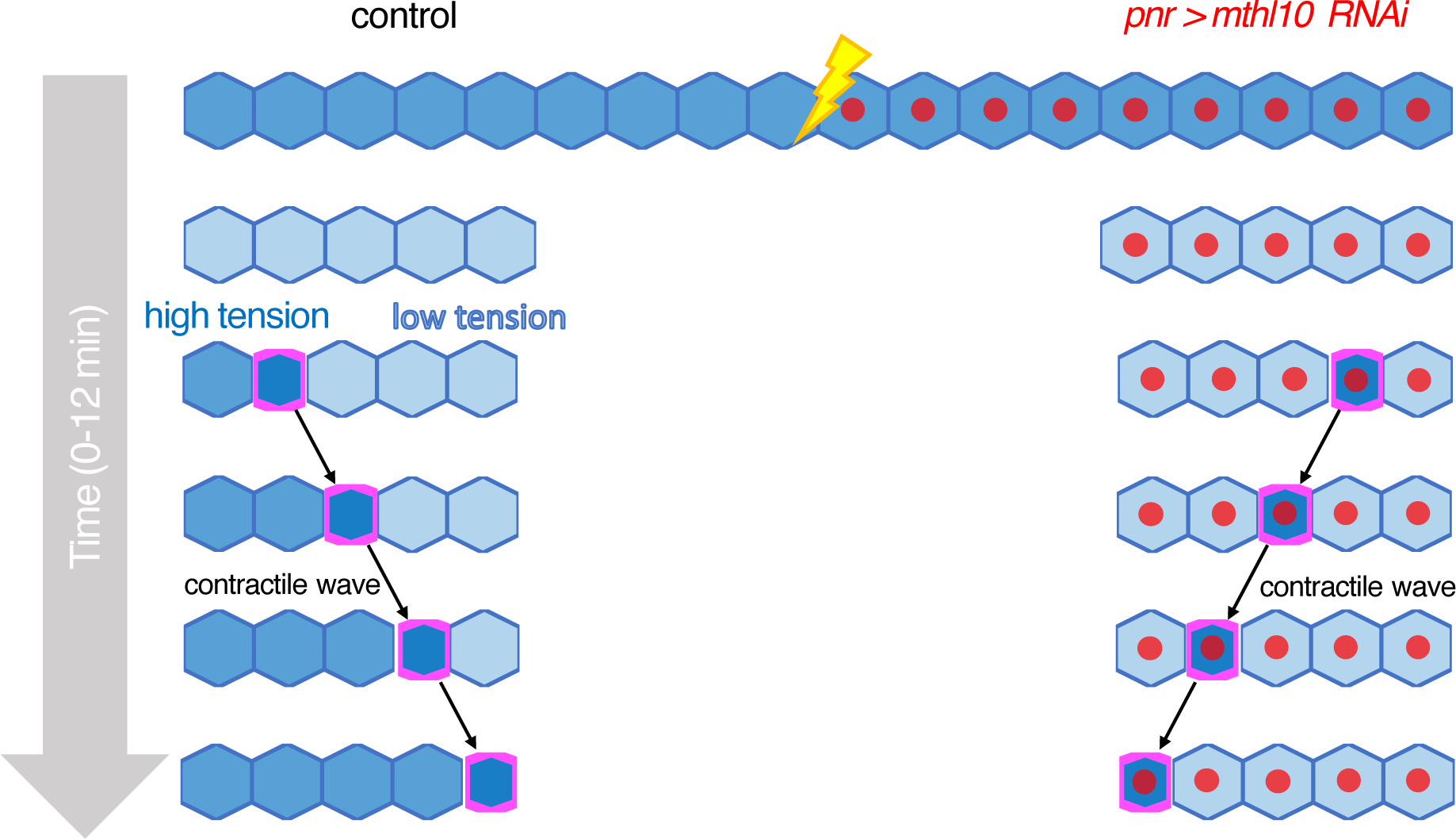
Schematic of how Mthl10-driven calcium signaling and the wound-induced contractile wave coordinate to restore tension around epithelial wounds.

Many systems that regulate cellular biomechanics have mechanical feedback, e.g., stretch-induced contractions (Odell *et al*., 1981; Zulueta-Coarasa and Fernandez-Gonzalez, 2018), that make the systems more responsive and robust. Is there any such feedback through Mthl10-regulated calcium signaling? Mthl10 is activated after wounding by the Gbp cytokines, which exist outside the cell in latent pro-Gbp forms and are converted to active cytokines by proteases released from the wound. A similar ligand-dependent Mthl10 activation occurs when isolated *Drosophila* wing discs are exposed to fly extract, which contains its own proteases and pro-Gbps. Interestingly, similar *in vitro* experiments demonstrate a mechanical sensitivity in extract-stimulated calcium signals (Narciso *et al*., 2017). It is not yet clear whether mechanical cues are involved in Mthl10 signaling *in vivo*, but several GPCRs are mechanically gated, and there are certainly mechanical changes upon wounding an epithelium that could modulate Mthl10 signaling.

In summary, this study reports an unexpected widespread reduction in tension around epithelial wounds. This loss of tension occurs within 30 seconds, and its restoration over 2-10 minutes is regulated by a cell surface receptor activating a signal transduction cascade, resulting in the release of cytoplasmic calcium. The receptor Mthl10 does not have an ortholog in vertebrates, but it is a member of the broadly conserved G_αq_-coupled GPCR family. It is possible that similar but not orthologous biochemical signaling cascades may be utilized by vertebrate epithelia to restore tension after wounding.

## MATERIALS AND METHODS

### Drosophila melanogaster genotypes

All fly genotypes used for recoil measurements had GFP-labeled E-cadherin (*P{Ubi-p63E-shg.GFP}5*, Kyoto DGGR 109007) in apical epithelial cell borders. They also had *UAS-mCherry.NLS* (BDSC 38424) – a monomeric red fluorescent protein with a nuclear localization signal – driven by *pnr-Gal4* (BDSC 25758) and temporally regulated by *tubP-Gal80^TS^* (BDSC 7017). The *pnr* domain is an hourglass-shaped domain in the center of the notum (Fig. 1C) that expresses *pannier (pnr)*, a gene required for dorsal closure (Herranz and Morata, 2001). The *Gal4-UAS* system allows us to drive genetic manipulations by placing the yeast enhancer *UAS* (Upstream Activation Sequence) upstream our gene of interest and having the yeast transcriptional activator protein Gal4 bind to *UAS* to activate the gene’s transcription (Brand and Perrimon, 1993). With *pnr-Gal4*, Gal4 is expressed only in the *pnr* domain, and thus genetic manipulations can be confined there, setting up a system where the area outside the *pnr* domain – called the control domain – can serve as an internal control. Gal80^TS^ is a temperature-sensitive protein that binds and inhibits Gal4 transcriptional activity at 18°C (Zeidler *et al*., 2004). At 29°C, however, Gal80^TS^ cannot bind Gal4 and can no longer repress it, activating Gal4-driven gene expression.

For tension mapping, the control genotype was *Ecad-GFP/+; pnr-Gal4, UAS-mCherry.NLS, TubP-Gal80^TS^/+*. Knockdown constructs were crossed into this line to be driven by *pnr-Gal4, tubP-Gal80^TS^*, to drive expression of *UAS-Rok RNAi* (VDRC 104675), *UAS-mthl10 RNAi* (BDSC 62315), or *UAS-IP_3_R RNAi* (BDSC 25937). The *mthl10 RNAi* knockdown results were verified with *UAS-mthl10 RNAi#2* (BDSC 51753).

For contractile wave tracking, cell shape changes were visualized with Ecad-GFP in the control fly genotype *Ecad-GFP/+; pnr-Gal4, UAS-mCherry.NLS, Gal80^TS^/+* and in the *mthl10* knockdown genotype *Ecad-GFP/UAS-mthl10 RNAi; pnr-Gal4, UAS-mCherry.NLS, tubP-Gal80^TS^/+.* The apical accumulation of myosin in contracting cells was visualized using the fluorescence signal of MyoII-GFP (BDSC 51564), a GFP-trapped endogenously tagged non-muscle myosin II heavy chain. Control genotype was *yw; MyoII-GFP/CyO* or *MyoII-GFP/+; pnr-Gal4, UAS-mCherry.NLS, Gal80^TS^/+*. Mthl10 knockdown genotype was *MyoII-GFP / UAS-mthl10-RNAi; pnr-Gal4, UAS-mCherry.NLS, Gal80^TS^/+*. The complete genotypes for each *Drosophila melanogaster* line used in this study are detailed in Table S1. The complete genotype for each figure panel and movie is provided in Table S2.

### Drosophila melanogaster husbandry

Flies were fed standard cornmeal-molasses food supplemented with dry yeast. *MyoII-GFP* flies without *Gal80^TS^* were maintained at room temperature (around 20-22°C). For control samples with *Gal80^TS^*, new vials were maintained at room temperature to inhibit Gal4 activation during embryogenesis and then incubated at 29°C for 2-5 days until experimentation to activate Gal4. For samples with *Rok RNAi, mthl10 RNAi,* or *IP_3_R RNAi* expressed in the *pnr* domain, animals were kept at 18°C for 3-5 days to inhibit Gal4 activation during embryogenesis and then moved to 29°C for 3-4 days until experimentation to activate Gal4.

### Pupal mounting

Animals were aged at 29° C for 12 to 18 hours after puparium formation (APF) or at room temperature for 15-23 hours APF and mounted with nota pressed against a coverslip (Shannon *et al*., 2017; O’Connor *et al*., 2022). The method is described in detail in O’Connor *et al*. (2022), but briefly, pupae were placed ventral side down onto double-sided tape (Scotch brand, catalog #665) on a microscope slide. The pupae were tilted about the anterior-posterior axis to allow access to the control domain. Each anterior puparium was removed using fine-tipped forceps to expose the notum (Fig. 1A). A 35 mm x 50 mm coverslip (Fisherbrand, cat#125485R) was prepared by layering two pieces of double-sided tape along one long edge. Then, the tape with dissected pupae was gently transferred to the coverslip such that the posterior tips of the pupae rested on the layers of double-sided tape on the coverslip and the nota were between the coverslip and piece of tape. This procedure allowed for nota to be pressed up against the coverslip for microscopy and laser ablation. An oxygen-permeable membrane (YSI, standard membrane kit, cat#1329882) was secured over the coverslip to prevent the pupae from becoming dehydrated or oxygen-deprived.

### Live imaging

*In vivo* live imaging was performed with a Nikon Ti2 Eclipse with X-light V2 spinning disc confocal microscope (Yokogawa CSU-X1 spinning disk head with Andor DU-897 EMCCD camera) using a 40X 1.3 NA oil-immersion objective. The camera and laser were positioned by moving ∼200 μm down the midline from the head-thorax joint and then laterally to the *pnr* border. For all samples, a pre-wound z-stack scan was taken using the 488-nm laser line to capture Ecad-GFP or MyoII-GFP and the 560-nm laser line to capture mCherry-labeled *pnr* nuclei. For tension measurements, 45-second videos were taken of Ecad-GFP labeled cell borders to track tricellular junction (TCJ) movement following micro-ablation. For contractile wave tracking, z-stacks were taken every 30 seconds after wounding or every minute when making before- and after-wave measurements. When imaging live, images at previous timepoints for the current movie can be viewed as soon as they are taken, allowing the contractile wave to be monitored almost in real-time in order to measure tension before and after the wave passed through (Fig. 3E-E′′, Movie 4).

### Laser wounding

All laser ablations for single-shot wounding and laser-induced recoil were performed with single pulses of the 3^rd^ harmonic (355 nm) of a Q-switched Nd:YAG laser (5 ns pulse-width, Continuum Minilite II, Santa Clara, CA). Laser pulse energy for the wound was adjusted to create a ∼70-μm radius of nuclear membrane damage, with specific energy values varying from 1.5 to 2.0 μJ. For scanned wounds, a series of micro-ablations with laser pulse energy of 0.4 μJ were created in the pattern of a circle with a radius of ∼40 μm, which matches the expected radius of cell lysis for a single-shot wound with a ∼70 μm radius of nuclear membrane damage (O’Connor *et al*., 2021a). The initial wound in both single-shot and scan-wounded samples always occurred on the *pnr* border, with the center of the wound positioned by moving ∼200 μm down the midline from the head-thorax joint and then moving laterally to the *pnr* border. Wounding on the *pnr* border allowed for symmetrical comparisons between *pnr* and control domain in internally controlled samples.

### Locations for tension measurements

Tension was measured in mediolateral (ML) and anterior-posterior (AP) borders at different radii lateral to the *pnr* border in both the *pnr* and control domains (Fig. 1B). The closest location in unwounded samples was 70 μm from the *pnr* border. The corresponding location in wounded samples was at the edge of the region of nuclear membrane damage as indicated by a disrupted mCherry signal in the *pnr* domain; a symmetric location was used on the control side. This radius ranged from 60-90 μm but was ∼70 μm in most samples. The second location was 110 μm from the *pnr* border in unwounded samples and 40 μm away from the edge of nuclear membrane damage in wounded samples. The farthest location was 210 μm from the *pnr* border in unwounded samples and 140 μm from the edge of nuclear membrane damage in wounded samples. For scan-wounded control samples, the location right outside of the scan, ∼40 μm from the *pnr* border, was tested for tension in addition to the 70, 110, and 210 μm locations. After initial tension mapping, ∼135 µm was added as a distance for tension testing to represent a single location distal to the wound. Unless otherwise specified, tension mapping was performed between 1 to 10 minutes post-wound in wounded samples and over a similar time in unwounded samples.

### Laser-induced recoil assay

Cortical tension, a combination of tension from the membrane, cytoskeleton, and cell adhesions, can be measured by laser-induced recoil. In this assay, an apical epithelial cell border labeled by Ecad-GFP is severed with a laser micro-ablation using the same laser that is used for wounding but at a lower power (Movie 1). Laser pulse energy for the laser-induced recoil micro-ablations was adjusted to ablate a single cell border, with specific energy values varying from 0.4 to 0.8 μJ. After ablating a cell border, the velocity at which the corresponding TCJs retract from each other is measured. In this assay, higher cortical tension leads to higher recoil velocities after ablation while lower cortical tension leads to lower recoil velocities after ablation (Fig. 1D′).

### Calculating recoil velocities

After ablating the cell border in laser-induced recoil, the distance between the corresponding TCJ’s of the severed border was measured every 2 seconds for 30 seconds with the length measurement tool in the Nikon NIS-Elements platform (RRID:SCR_014329) or ImageJ (Schindelin *et al*., 2012). TCJ’s show up as brighter spots of Ecad-GFP fluorescence. The first frame with visible TCJ’s after micro-ablation was set as t = 0 s. Based on a sampling of 60 experiments, this first TCJ-visible frame occurred at 0.93 ± 0.28 s. For each sample, the length measurements were analyzed in Mathematica (Wolfram Research Inc., Champaign, IL) to calculate the average recoil velocity (in nm/s) over the first 30 seconds based on a nonlinear model, *d*(*t*) = *d*_0_ + *d*_1_(1 – *e^−t/τ^*), where *d*(*t*) is distance between TCJ’s at time *t* after micro-ablation and *d_0_*, *d_1_* and *^τ^* are fitting parameters. Six example data sets and fits are shown in Fig. S5A-F. Note that the recoils are quite small, typically just 1-2 µm over 30 s. Given these small recoil extents, we chose to increase the signal-to-noise ratio of our measurements by using the average recoil velocity over 30 s, <*v*>_0-30_ rather than estimating the instantaneous recoil velocity at time zero, *v*_0_. A comparison of these two measures is shown in Fig. S5G for all 828 recoil experiments reported here (across all locations, edge orientations, and genotypes). For this complete data set, the median value of v_0_ was 70.9 nm/s and the median value of v_0_’s standard error was 13.0 nm/s. In comparison, the median value of <v>_0-30_ was 38.3 nm/s and the median value of its standard error was just 2.2 nm/s. Thus, even though estimates of <v>_0-30_ were about half v_0_, they had standard errors that were just 1/6 as large, which translates into an increase in the signal-to-noise ratio by a factor of ∼3.

An important caveat to using such temporal averaging is that <v>_0-30_ is dependent not just on the local cell-edge tension and viscous damping coefficient, but also on the effective spring constant through its impact on how the pulling forces from adjacent cell edges change as the TCJs move. This effective spring constant is expected to depend on the local triple-junction angles and the cell-edge tension in the adjacent edges. Measurements of cortical tension based on such temporal averaging of the recoil velocity should thus be considered a weighted average of cortical tension in the ablated edge and its neighbors.

### Cell contraction analysis

The contractile wave can be visualized over time using Ecad-GFP, which labels apical epithelial cell borders (Movies 3, 4, and 5). To quantify these contractions, we calculated the vector displacement field 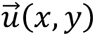 between successive images (taken 30 s apart) using Mathematica’s ImageDisplacements function. The displacement fields were then used to calculate the local area strain 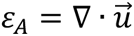 by integrating the displacements over the boundary of a 7×7 square centered on each pixel and applying the divergence theorem: 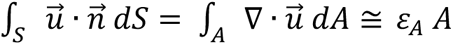. The size of the square used for integration was a compromise that increased the signal-to-noise ratio without over-smoothing the strain fields. For visualization, the strain field was color coded using a TemperatureMap scheme and overlaid on an inverted gray-scale image of the Ecad-GFP channel. All strain calculations and visualizations were performed in Mathematica (Wolfram Research Inc., Champaign, IL).

### MyoII-GFP intensity analysis

To quantify MyoII-GFP intensity as the contractile wave travels inwards, we used the ImageJ plug-in Radial Profile Extended (Ferri *et al*., 2008) to plot the average MyoII-GFP intensity profile as a function of distance from the center of the wound. For each MyoII-GFP movie, radial profile angle analysis was applied to images every 2 minutes starting from 2 minutes post-wound and ending at 14 minutes post-wound (Fig. 3B). A custom R script (R Core Team, 2021) was then used to determine each timepoint’s peak MyoII-GFP intensity and its distance from the wound. This distance was used to represent the wave’s location for calculating the average distance of the wave from the wound over time, which was graphed in Microsoft Excel (Fig. 3C).

### Statistical analysis

All statistical analyses were performed with GraphPad Prism version 9.2.0, which was also used to graph the recoil velocity values. Each graph displays the value for each sample as a point, with bars representing mean and SEM. Data from the control domain are indicated by negative distance values and shown in black. Data from the *pnr* domain are indicated by positive distance values and shown in red when genetically manipulated. Statistical analyses were conducted by one-way ANOVA with multiple comparisons for recoil velocities from each genotype. For unwounded control and *mthl10 RNAi* samples, along with the Fig. 2 graphs showing post-wound tension over time and Fig. S2B, the mean of each column was compared with the mean of every other column. For unwounded *Rok RNAi* and the rest of the wounded samples, means were compared between the most proximal and distal locations on each domain, and then between control and *pnr* domains for the most proximal and distal locations. For scan-wounded samples, the 40 µm location was left out of statistical tests because this location was not tested in other types of samples. For Fig. 3 and Fig. 4H, paired sample t-tests were used to compare the two groups, with each sample producing one pair of tension measurements. For Fig. 4G, an unpaired t-test was used to compare the two groups.

### Vertex model

Simulations of vertex models were run in Mathematica (Wolfram Research Inc., Champaign, IL) following methods and parameter choices similar to those used previously to model wound healing in Drosophila wing imaginal discs (Tetley *et al*., 2019). Briefly, the tissue is modeled as a set of fully connected polygonal cells and the mechanical energy (*E*) of the entire tissue is given by:

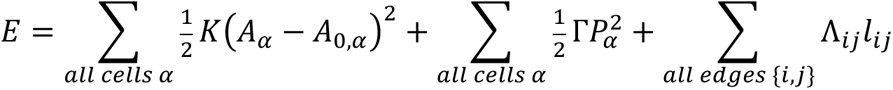

where the first sum represents the energy required to stretch or compress each cell away from its target area *A*_0,*α*_; the second sum represents cortical contractile energy proportional to the squared perimeter of each cell 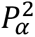; and the third sum represents a combination of cortical tension and cell-cell adhesion that yields a line tension Λ*_ij_* along the length of each polygonal edge, *l_ij_*. The parameters *K* and Γ are the area elasticity modulus and cortical contractility, respectively. In a simulation, the tissue geometry evolves over time according to the forces exerted on each vertex *F_i_* = − ∂*E*⁄∂*x_i_* and overdamped dynamics, ∂*x_i_*⁄∂*t* = *F_i_*⁄μ, where μ is the damping coefficient. Parameters for the vertex model were as follows: normalized contractility Γ⁄*KA*_0_ = 0.01771 for normal cells and 0.00664 for those with reduced contractility; average normalized line tension 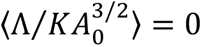 with fluctuations of standard deviation σ_m_ = 0.01 and tension recovery time τ_m_ = 25 s to model myosin II fluctuations (Curran *et al*., 2017; Tetley *et al*., 2019); and friction coefficient µ = 30 s. An isotropic, far-field tensile stress Σ was implemented as an additional force acting normal to the outer edge of the patch with normalized strength Σ⁄*KA*_0_ = 0.14. This value was chosen so that releasing this far-field stress would cause the cell patch to shrink by 15%, as observed when a large circular patch of the mid- to late-stage pupal notum (18-24 hours APF) is separated from the surrounding tissue by laser ablation (Bonnet *et al*., 2012). Cell neighbor exchanges were implemented by monitoring the geometry at each time step and applying a T1 transition for any edges that were shorter than 0.07% of the mean edge length.

A circular patch of 225 cells was initialized by Voronoi tessellation of a circular area with randomly distributed seed points and then equilibrated by running the model without stochastic fluctuations until the patch reached a steady state. To simulate recoil-based tension measurements, we ran the equilibrated model forward with stochastic fluctuations and calculated the force a vertex would experience if one of its adjacent edges were ablated – i.e., setting the edge’s line tension and its adjacent cells’ contractility and area elasticity all to zero.

## Supporting information

Recoil velocity data

## ACKNOWLEDGEMENTS

Special thanks to Jasmine Su for technical assistance with measuring tension within a minute after wounding. Thank you to Dr. Indrayani Waghmare, Aubrie Stricker, Kimi LaFever Hodge, Elkie Peebles, Dr. Aaron Stevens, Mia Grace Cantrell, Gemini Simpson, and Jasmine Su for their mentorship, feedback, and support. We would also like to thank Dr. Shigeo Hayashi, the Bloomington *Drosophila* Stock Center (BDSC), Vienna *Drosophila* Resource Center (VDRC), and the Kyoto Department of *Drosophila* Genomics and Genetic Resources (Kyoto DGGR) for *Drosophila* lines. Funding: This work was supported by the National Institute of General Medical Sciences 1R01GM130130 and by Vanderbilt Seeding Success to both A.P.M and M.S.H; by the Harold Sterling Vanderbilt Honors Scholarship to I.S.H.; and by the American Heart Association (19PRE34410069) to J.T.O.

## SUPPLEMENTAL MATERIALS

**Figure S1.**
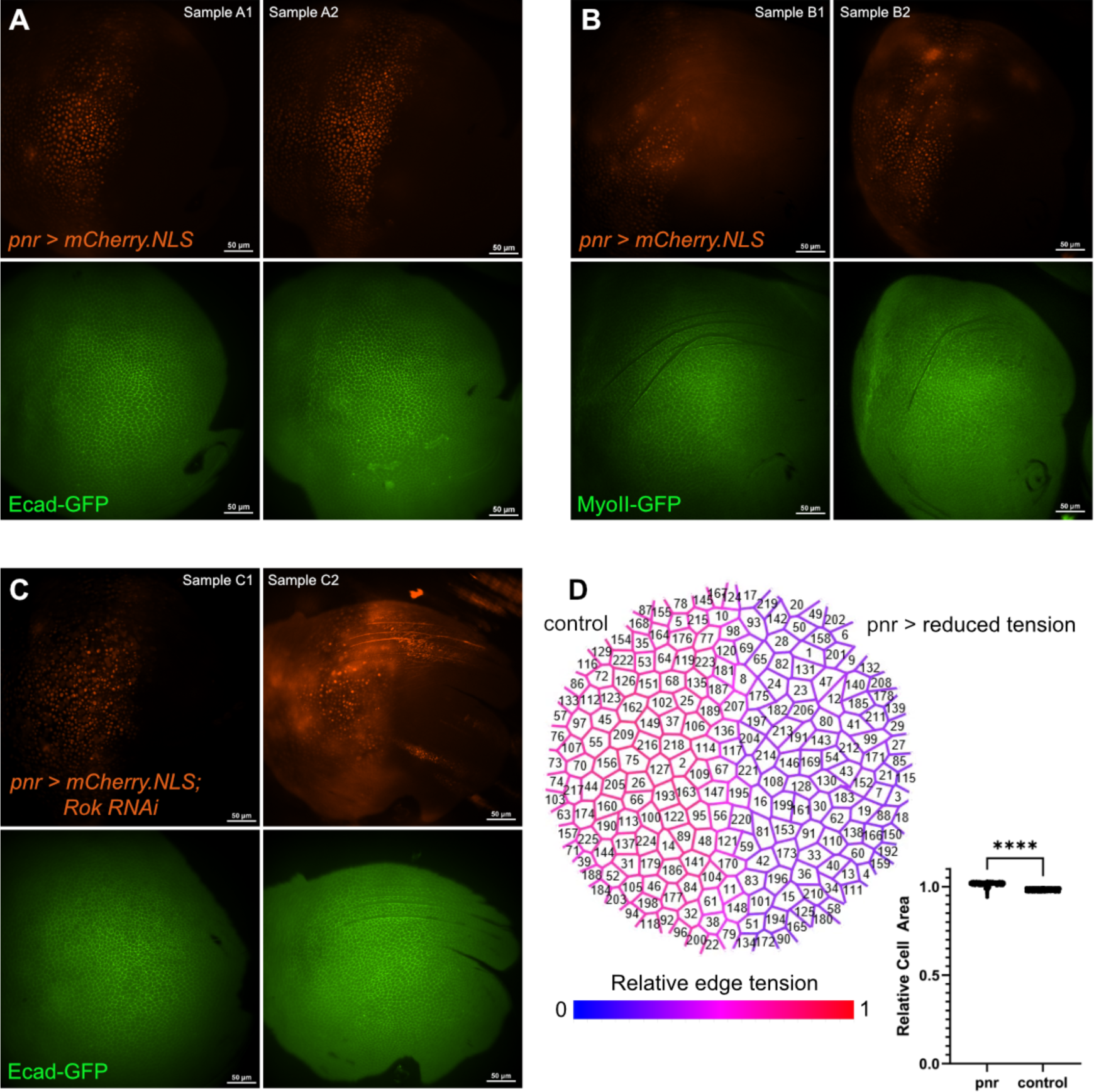
Cell morphology in the *pnr* and control domains in the *Drosophila* pupal notum. **(A-B)** Two samples each in which mCherry.NLS is expressed only in the *pnr* region (top) while Ecad-GFP or MyoII-GFP is expressed throughout the notum (bottom). In each case, the adjacent *pnr* and control domains have no discernable difference in cell morphology or the brightness levels of Ecad- or MyoII-GFP. The few dark lines are folds in the tissue that sometimes occur during mounting. **(C)** Two samples in which mCherry.NLS and *Rok-RNAi* are expressed only in the *pnr* region (top) while Ecad-GFP is expressed throughout the notum (bottom). Even with this *pnr*-specific knockdown of Rok, the adjacent regions still have no discernable difference in cell morphology or Ecad-GFP brightness levels. **(D)** Results of an equilibrated vertex model that simulates a *pnr*-specific knockdown of Rok by locally reducing contractility; relative cell-edge tension is reduced to approximately half (59%). Such models automatically compensate by allowing cell areas in the *pnr* region to increase further from their target areas, thus increasing tension in the model’s area-based terms. The increase in cell area in the model is statistically significant, but small (3%). Such a small increase would be dwarfed in experiments by the natural variance in cell areas.

**Figure S2.**
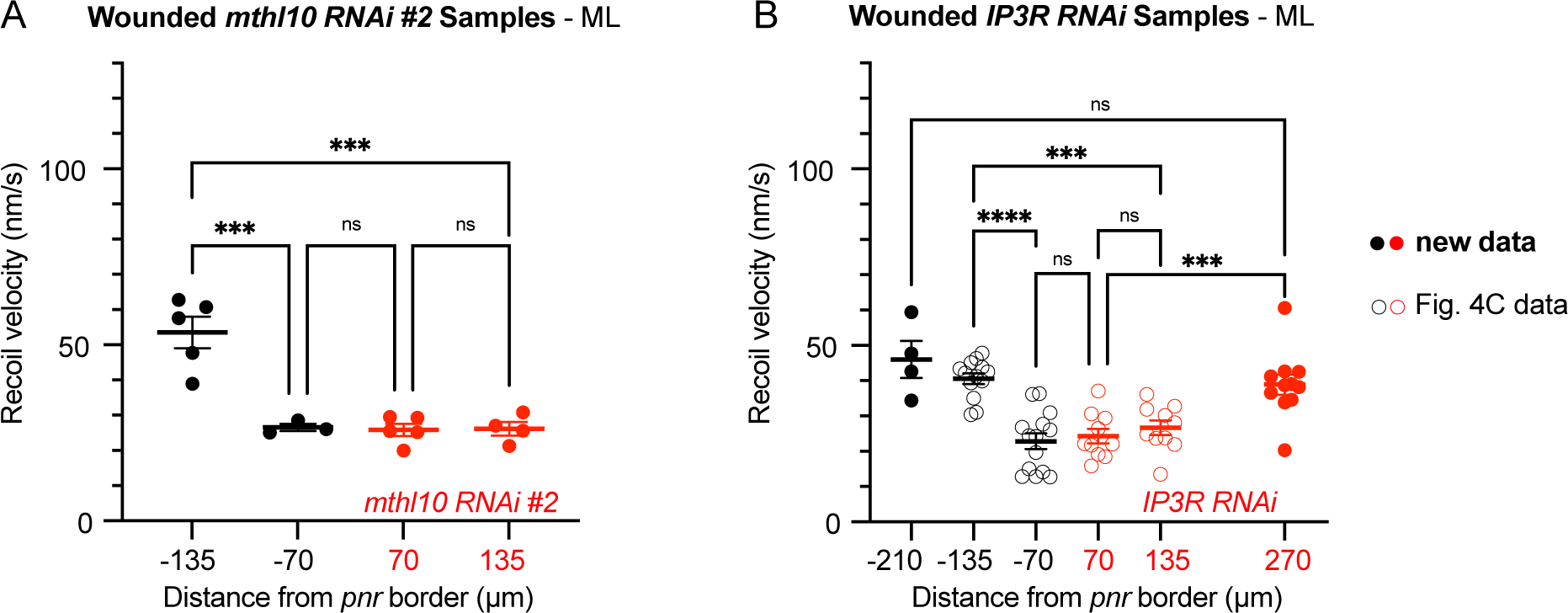
Additional tension mapping with *pnr > mthl10 RNAi #2* and *pnr > IP3R RNAi*. **(A)** In wounded *pnr > mthl10RNAi #2* samples, tension loss persisted in the knockdown region in measurements taken 1-10 minutes post-wound, verifying the first *mthl10 RNAi* line; *n* = 5, 3, 5, and 4 **(B)** Tension was reduced at ∼70 µm and ∼135 µm in the knockdown region for *pnr > IP3R RNAi* but not at ∼270 µm, suggested that IP3R is required to restore wound-induced tension loss but this initial tension loss is confined to a limited region that does not extend to ∼270 µm from the wound. These measurements were taken 1-10 minutes post-wound. Graph bars represent mean ± SEM; *n* = 4, 13, 14, 10, 10 and 11; ***p<0.001, ****p<0.0001 by one-way ANOVA with multiple comparisons.

**Figure S3.**
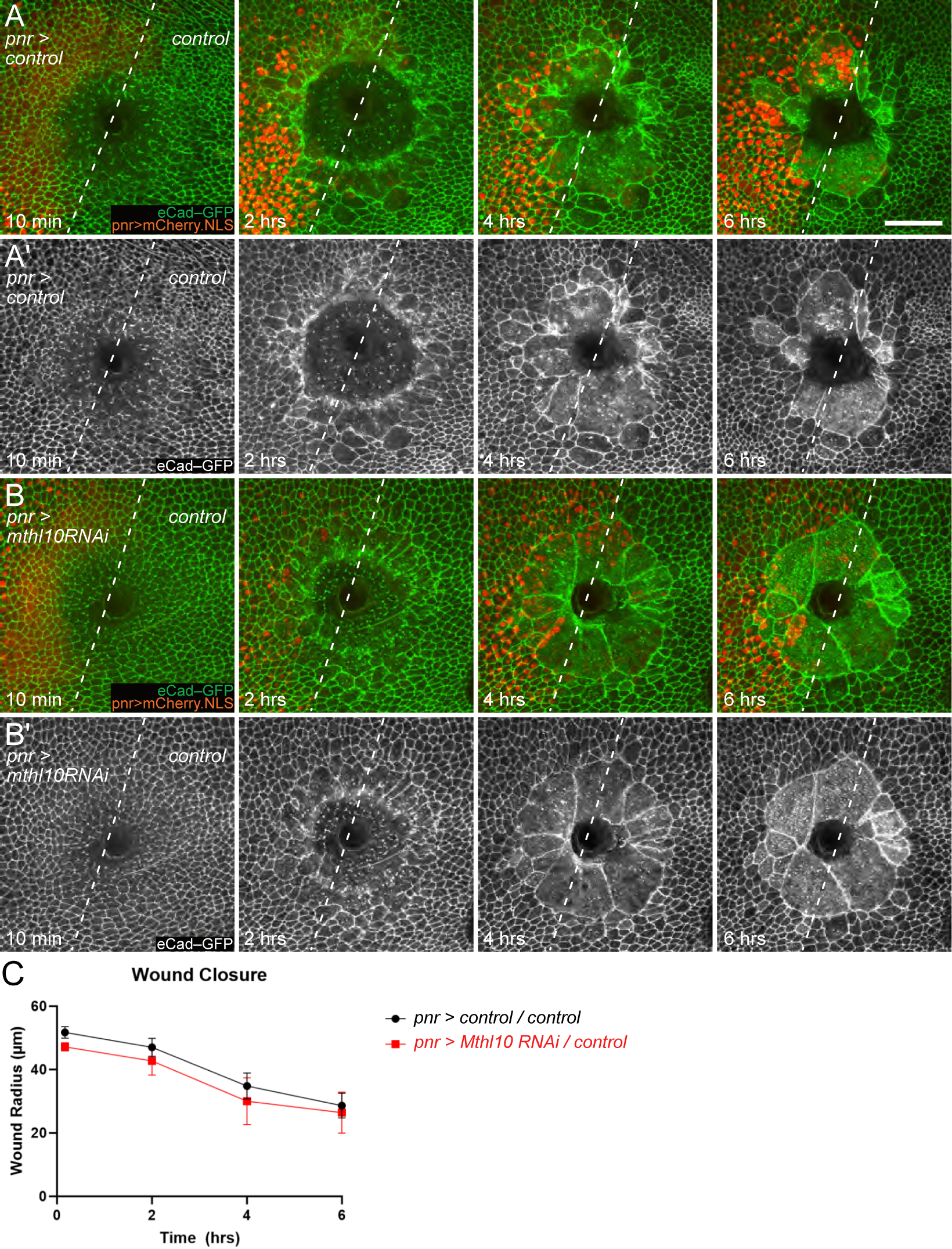
Knocking down Mthl10 does not impact the rate or timing of wound closure. (A, A’) Control pupal wound, injured on the border of the *pnr-Gal4* expression domain, repairs over the course of ∼6 hours. Syncytia are evident as the wound closes. Closure is obscured by a melanized scab over the wound; *n* = 5. **(B)** When *Mthl10-RNAi* is expressed in the *pnr* domain, both sides repair at approximately the same rate as in control pupae; *n* = 4. There are also no clear differences in syncytia formation or cellular migration towards the wound site between cells expressing *Mthl10-RNAi* and internal control domain cells. Scale bar = 50 µm. **(C)** Quantification of wound radius as a function of time shows no difference in wound closure rate when comparing *pnr > control/control* wounds versus *pnr > Mthl10 RNAi/control* wounds.

**Figure S4.**
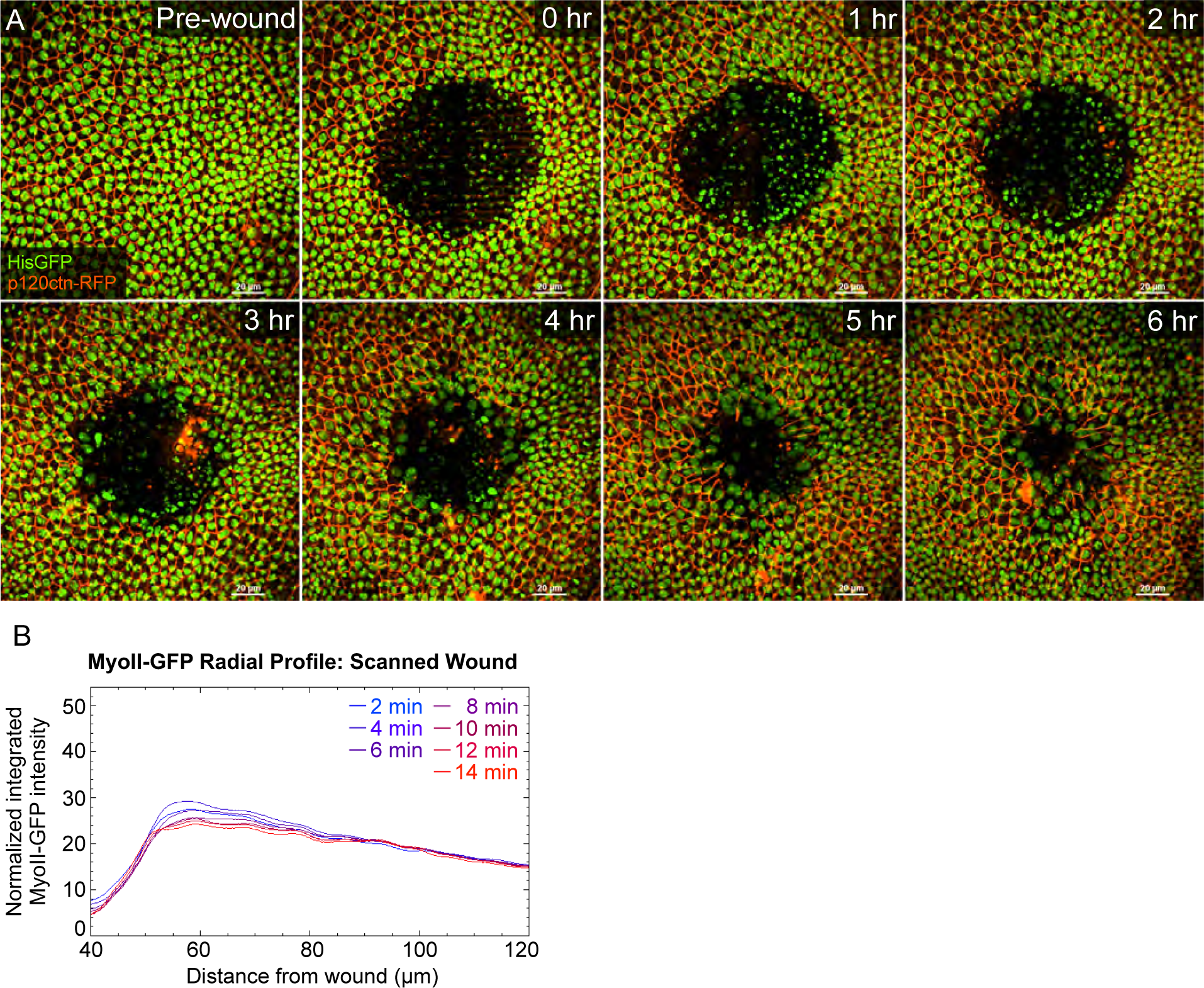
Scanning Ablation. **(A) Lower-power scanning ablation kills exposed cells with minimal damage to surrounding cells.** Time-lapse images of wounds after scanning ablation of a pupal notum expressing both p120ctn-RFP to mark cell borders and His2AvGFP to label chromatin. Scanning ablation yields an exposed region with lots of chromatin fragments that are immobile and still fluorescently labeled. These fragments are cleared as the wound is closed over the next 6 hours by cells moving into the wound bed from outside the exposed region. Scale bar = 20 µm. **(B)** Scanning ablation yields no apparent contractile wave. Radial profile analysis of MyoII-GFP fluorescence around a scanned wound, based on the images shown in Figure 5 and Movie 8. Note the lack of an inward-moving peak of MyoII-GFP fluorescence.

**Figure S5.**
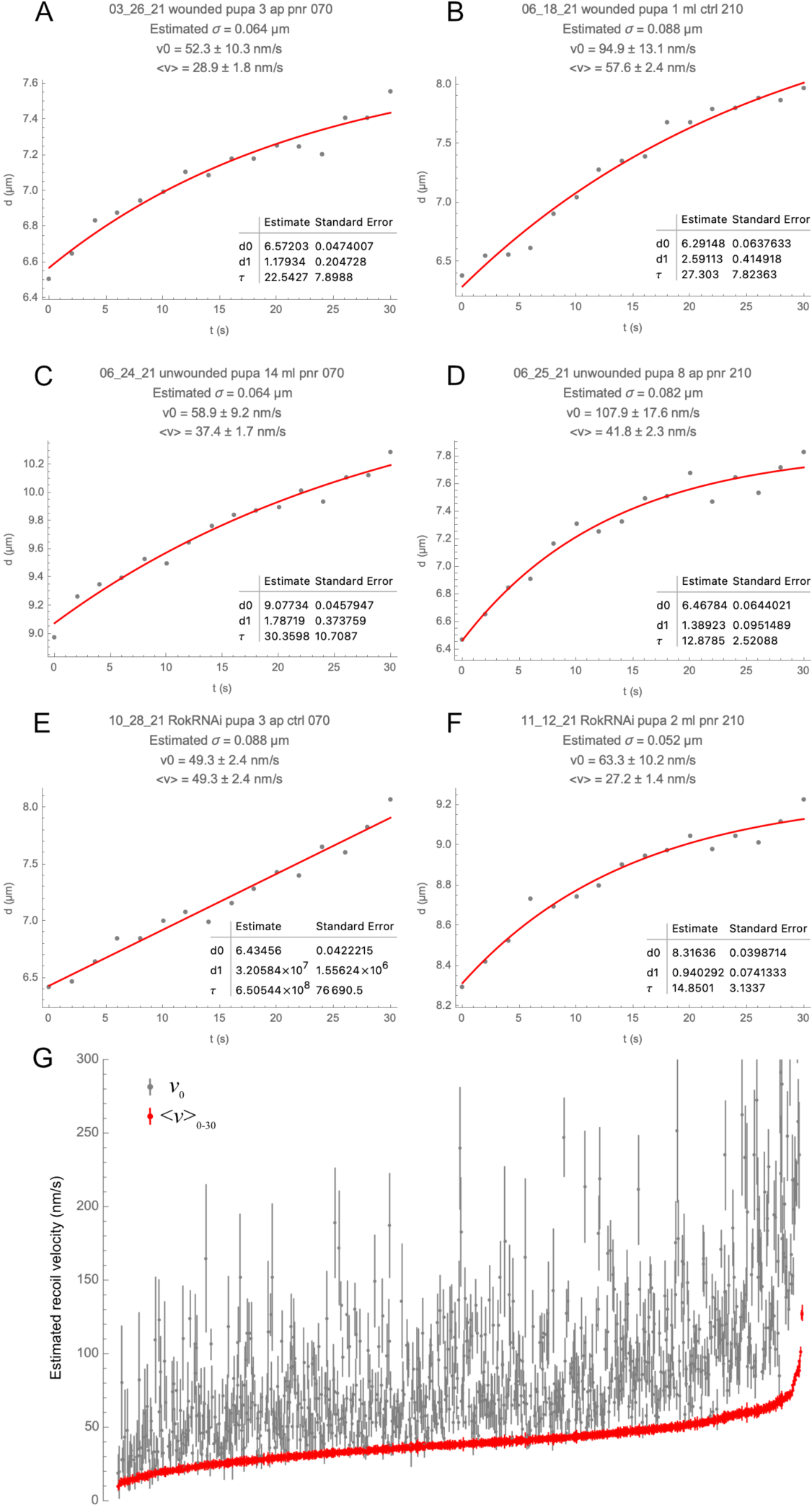
Estimating recoil velocities from post-ablation displacement data. **(A-F)** Six example data sets with nonlinear model fits overlain on data points. The heading above each plot notes the genotype, wounding condition, estimated uncertainty in position estimate (σ), and recoil velocities v_0_ and <v>_0-30_ respectively estimated from the slope at *t* = 0 or the average velocity over 30 s. **(G)** Comparison of v_0_ and <v>_0-30_ estimates across 833 measurements over all genotypes, wounding conditions, and distances from the wound. Each point represents an estimate ± its standard error. Paired v_0_ and <v>_0-30_ estimates are aligned horizontally. Note that estimates of <v>_0-30_ are about half as large as estimates of v_0_, but standard errors of <v>_0-30_ estimates are just 1/6 those of v_0_. Using <v>_0-30_ rather than v_0_ thus increases the signal-to-noise ratio by a factor of ∼3.

**Table S1.** Identity of Drosophila genetic elements used in this study.

| **Fly genetic element** | **Complete genotype** |
| --- | --- |
| “*Ecad-GFP*” | *w*; {*ubi-p63E-shg.GFP*, *w+*}5<br>(FlyBase ID: FBtp0014096) |
| “*MyoII-GFP*” | *w*[*]; *P*{*w*[+*mC*]=*PTT-GC*}*zip*[*CC01626*]/*SM6a*<br>(FlyBase ID: FBst0051564) |
| “*pnr-Gal4*” | *P*{*w*[+*mC*]=*UAS-Dcr-2.D*}1, *w*[1118];<br>*P*{*w*[+*mW.hs*]=*GawB*}*pnr*[*MD237*]/*TM3*, *Ser*[1]<br>(FlyBase ID: FBst0025758) |
| “*tubP-Gal80<sup>ts</sup>*” | *w*[*]; *P*{*w*[+*mC*]=*tubP-GAL80[ts]*}2/*TM2*<br>(FlyBase ID: FBst0007017) |
| “*UAS-IP3R RNAi*” | *y*[1] *v*[1]; *P*{*y*[+*t7.7*] *v*[+*t1.8*]=*TRiP.JF01957*}*attP2*<br>(FlyBase ID: FBst0025937) |
| “*UAS-mCherry.NLS*” | *w*[*]; *P*{*w*[+*mC*]=*UAS-mCherry.NLS*}3<br>(FlyBase ID: FBst0038424) |
| “*UAS-mthl10-RNAi*” | *y*[1] *v*[1]; *P*{*y*[+*t7.7*] *v*[+*t1.8*]=*TRiP.HMJ23672*}*attP40/CyO*<br>(FlyBase ID: FBst0062315) |
| “*UAS-mthl10-RNAi#2*” | *y*[1] *v*[1]; *P*{*y*[+*t7.7*] *v*[+*t1.8*]=*TRiP.HMC03304*}*attP2*<br>(FlyBase ID: FBst0051753) |
| “*UAS-Rok-RNAi*” | *P*{*KK107802*}*VIE-260B*<br>(FlyBase ID: FBst0476533) |
| “*HisGFP*” | *w*[*]; *P*{*w*[+*mC*]=*His2Av-EGFP.C*}2/*SM6a*<br>(FlyBase ID: FBti0077861) |
| “*p120ctn-RFP*” | *yw*; *M*{*sqh-p120ctn.TagRFP*} (on 2<sup>nd</sup> chromosome)<br>(FlyBase ID: FBal0344828)<br>from Shigeo Hayashi |

**Table S2.**
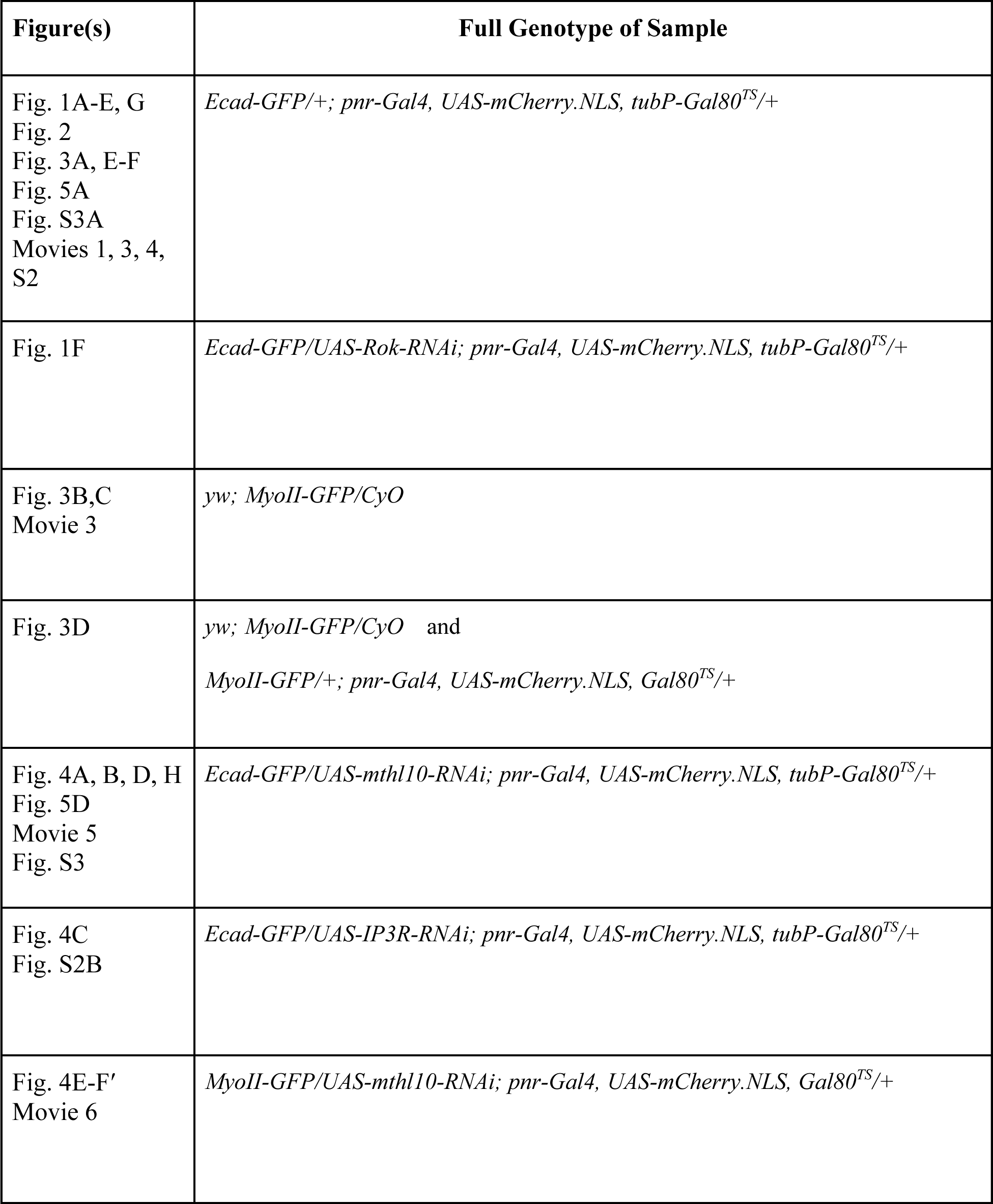
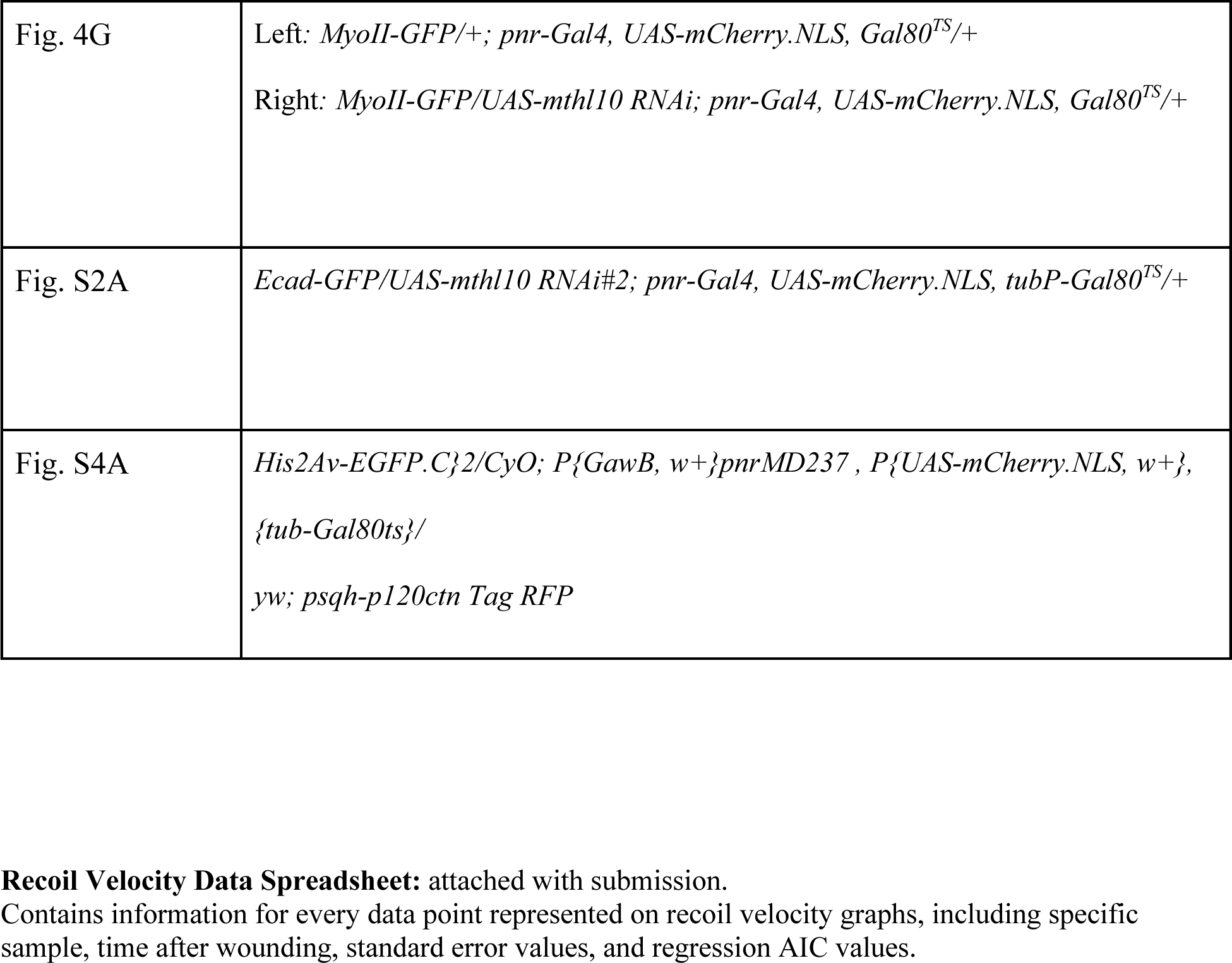
Genotypes of flies in each figure panel.

| **Figure(s)** | **Full Genotype of Sample** |
| --- | --- |
| Fig. 1A-E, G<br>Fig. 2<br>Fig. 3A, E-F<br>Fig. 5A<br>Fig. S3A<br>Movies 1, 3, 4,<br>S2 | *Ecad-GFP/+; pnr-Gal4, UAS-mCherry.NLS, tubP-Gal80<sup>TS</sup>/+* |
| Fig. 1F | *Ecad-GFP/UAS-Rok-RNAi; pnr-Gal4, UAS-mCherry.NLS, tubP-Gal80<sup>TS</sup>/+* |
| Fig. 3B,C<br>Movie 3 | *yw; MyoII-GFP/CyO* |
| Fig. 3D | *yw; MyoII-GFP/CyO* and<br>*MyoII-GFP/+; pnr-Gal4, UAS-mCherry.NLS, Gal80<sup>TS</sup>/+* |
| Fig. 4A, B, D, H<br>Fig. 5D<br>Movie 5<br>Fig. S3 | *Ecad-GFP/UAS-mthl10-RNAi; pnr-Gal4, UAS-mCherry.NLS, tubP-Gal80<sup>TS</sup>/+* |
| Fig. 4C<br>Fig. S2B | *Ecad-GFP/UAS-IP3R-RNAi; pnr-Gal4, UAS-mCherry.NLS, tubP-Gal80<sup>TS</sup>/+* |
| Fig. 4E-F'<br>Movie 6 | *MyoII-GFP/UAS-mthl10-RNAi; pnr-Gal4, UAS-mCherry.NLS, Gal80<sup>TS</sup>/+* |
| Fig. 4G | Left: *MyoII-GFP/+; pnr-Gal4, UAS-mCherry.NLS, Gal80<sup>TS</sup>/+*<br>Right: *MyoII-GFP/UAS-mthl10 RNAi; pnr-Gal4, UAS-mCherry.NLS, Gal80<sup>TS</sup>/+* |
| Fig. S2A | *Ecad-GFP/UAS-mthl10 RNAi#2; pnr-Gal4, UAS-mCherry.NLS, tubP-Gal80<sup>TS</sup>/+* |
| Fig. S4A | *His2Av-EGFP.C*{2/CyO; *P*{*GawB, w+*}*pnrMD237*, *P*{*UAS-mCherry.NLS, w+*},<br>*{tub-Gal80ts}*/<br>*yw; psqh-p120ctn Tag RFP* |

**Supplemental Movies 1-8:** attached with submission.

**Movie 1. Laser-induced recoil example**

Tricellular junctions of an ablated cell border retract after laser microablation. Quantifying the recoil velocity of retraction was used to estimate cortical tension. Movie shows a control sample mediolateral border being ablated and then recoiling for ∼60 seconds. The bright dots in the 3^rd^ frame are a stroboscopic image of the spinning disk’s pinhole pattern generated by a flash of white-light emission from plasma recombination during laser ablation. Scale bar = 10 µm.

**Movie 2. Contractile wave visualized by cell contractions in wounded control sample**

The post-wound contractile wave was visualized with Ecad-GFP labeling apical epithelial cell borders as they contract in concentric rings one at a time, moving closer to the wound over time. The first half of the movie shows a control sample from 0-15 minutes every 30 seconds after wounding. The second half shows the same sample but with an inverted Ecad-GFP signal and an overlay of the calculated area strain between successive frames. Scale bar = 50 µm.

**Movie 3. Contractile wave visualized by myosin accumulation in wounded control sample**

The post-wound contractile wave was visualized with MyoII-GFP labeling non-muscle myosin II, which temporarily accumulates apically in each cell before it contracts. Movie shows a control sample from 0-15 minutes every 30 seconds after wounding. Scale bar = 50 µm.

**Movie 4. Monitoring contractile wave to measure tension before and after**

Tension was measured before and after the contractile wave passed through by using Ecad-GFP to monitor the wave. Movie shows a control sample from 4-10 minutes every minute after wounding, right after a before-wave tension measurement is taken and right before an after-wave tension measurement is taken. Scale bar = 50 µm.

**Movie 5. The contractile wave is not disrupted in *pnr > mthl10-RNAi* samples (Ecad-GFP)**

The contractile wave, visualized with Ecad-GFP, was not disrupted in *pnr > mthl10-RNAi* samples. *mthl10* is knocked down on the left side of the wound. The first half of the movie shows a *pnr > mthl10 RNAi* sample with images taken every 30 seconds from 0-14 minutes after wounding, and with a pre-wound image containing the red channel (*mCherry.NLS*) to indicate the boundaries of the *pnr* knockdown domain. The second half shows the same sample but with an inverted Ecad-GFP signal and an overlay of the calculated area strain between successive frames. Scale bar = 50 µm.

**Movie 6. The contractile wave is not disrupted in *pnr > mthl10-RNAi* samples (MyoII-GFP)**

The contractile wave, visualized with MyoII-GFP, was not disrupted in *pnr > mthl10-RNAi* samples. *mthl10* is knocked down on the left side of the wound. Movie shows a *pnr > mthl10-RNAi* sample with images taken every 30 seconds from 0-16 minutes after wounding, and with a pre-wound image containing the red channel (*mCherry.NLS*) to indicate where the *pnr* knockdown domain is. Scale bar = 50 µm.

**Movie 7. No apparent contractile wave after scan-wounding (Ecad-GFP)**

With Ecad-GFP labeling cell borders, there was no evidence of a contractile wave around scanned wounds. The first half of the movie shows a control sample from 2-15.5 minutes every 30 seconds after scan-wounding. The second half shows the same sample but with an inverted Ecad-GFP signal and an overlay of the calculated area strain between successive frames. Scale bar = 50 µm.

**Movie 8. No apparent contractile wave after scan-wounding (MyoII-GFP)**

With MyoII-GFP labeling non-muscle myosin II, there was no evidence of a contractile wave around scanned wounds. Movie shows a control sample from 2-15.5 minutes every 30 seconds after scan-wounding. Scale bar = 50 µm.

## Notes

### Competing Interest Statement

The authors have declared no competing interest.

### Summary of Updates

Corrected an error in the schematic in Figure 6.

